# DAP12-deficient microglia adopt a TREM2-independent antigen-presenting state with a glial-to-neuronal cascade in Nasu-Hakola disease

**DOI:** 10.64898/2026.10.07.757387

**Authors:** Julio Leon, Satoshi Yoshinaga, Ikuko Koya, Grecia Palafox, Yoshinari Ando, Ken-ichiro Kubo, Jonathan Moody, Masaki Takao, Piero Carninci, Jay Shin, Carlo Condello, Hideyuki Okano, Chung Chau Hon, Satoru Morimoto

**Author notes:** **Corresponding authors.** Julio Leon and Satoru Morimoto.

## Abstract

Nasu-Hakola disease (NHD), a dementia with about 200 cases worldwide, is caused by loss-of-function mutation of *TYROBP*, encoding DAP12, or its receptor *TREM2*. TREM2-DAP12 is the switch that converts microglia from a surveillant to a reactive state; what DAP12-deficient microglia become and to what extent they reshape their microenvironment is unknown. We profiled prefrontal cortex and hippocampus from an NHD donor and control by single-nucleus RNA sequencing and mapped the glial changes *in situ* by spatial transcriptomics. NHD microglia adopted a CIITA-driven, MHC class II-high antigen-presenting state with complement, iron and glycolytic programs and Fc receptors but not the TREM2-dependent lipid program. Astrocytes became pan-reactive and lost metallothionein expression, oligodendrocytes lost mature cells and cholesterol synthesis, and excitatory neurons lost presynaptic and activity programs, consistently between regions and across a published cohort. DAP12 deficiency left microglia in an antigen-presenting state short of the TREM2-dependent arm, accompanied by a glial-to-neuronal cascade.

## Introduction

Nasu-Hakola disease (NHD; polycystic lipomembranous osteodysplasia with sclerosing leukoencephalopathy) is a recessive early-onset dementia caused by biallelic mutation of *TYROBP*, which encodes the signaling adaptor DAP12, or of its partner receptor *TREM2*^1,2^. Patients present in the fourth to fifth decade with bone cysts and fractures, a progressive frontal syndrome, cerebral atrophy, basal-ganglia calcification and sclerosing white-matter degeneration^3–5^. Most disease-causing mutations abolish or reduce the protein^6,7^, and mutation of either subunit produces the same brain disease^2^. Approximately 200 cases have been reported worldwide^8^, and genetically confirmed postmortem NHD brain is exceptionally scarce.

DAP12 is a hub of immunoreceptor tyrosine-based activation motif (ITAM) signaling in tissue macrophages. It couples TREM2, TREM1, SIRPβ1, CLEC5A and the activating CD300 receptors to the kinase SYK^9^, and it is required for CSF1R-driven macrophage proliferation and survival^10^. With the related adaptor FcRγ it also drives bone resorption by osteoclasts^11^, which is why NHD affects bone as severely as brain and why *TREM2* mutations, which spare the other DAP12-coupled receptors, can present as dementia without bone cysts^12^. Loss of DAP12 therefore abolishes signaling through several receptors at once, whereas *TREM2* variants that raise the risk of Alzheimer’s disease remove one and do not cause NHD^13^; monoallelic loss of *TYROBP* itself was recently reported to raise that risk^14^. The same genetics places microglia at the center of the common disease, since genome-wide association studies map much of late-onset Alzheimer’s-disease risk to microglial genes such as *TREM2*, *CD33* and *PLCG2*^15^; what a microglial cell does when one node of that network fails completely is therefore a question the common disease shares with NHD.

In the brain, both subunits are expressed by microglia, which regulate astrocytes, oligodendrocytes, and synapses through what they secrete and remove^16–18^, and the TREM2-DAP12 complex is the switch that converts them from a surveillant to a reactive state. In mouse models of neurodegeneration, that conversion follows a two-step trajectory into a disease-associated microglial (DAM) state, also termed the microglial neurodegenerative phenotype (MGnD): a first, TREM2-independent step that lowers homeostatic checkpoints such as *P2RY12*, and a second, TREM2-dependent step that installs a lipid-handling and phagocytic program (*LPL*, *CST7*, *ITGAX*, *CLEC7A*)^19,20^. Because only the second step requires the complex, we expect NHD microglia not to progress to the TREM2-dependent state; what they become instead, and to what extent that state alters their microenvironment, remain unknown.

The human evidence is limited and cannot be supplied by models. *Tyrobp*- or *Trem2*-deficient mice show hypomyelination, synaptic degeneration, and impaired clearance of myelin debris, but only mild microglial changes that do not recapitulate the human disease^17,21,22^. NHD microglia have been described by immunohistochemistry as mildly activated in white matter with little or no DAP12 protein^7,23^, and TREM2-null microglia derived from patient stem cells show lysosomal dysfunction with loss of cholesterol and lipid-droplet genes^24^. The single-nucleus study by Zhou et al., which focused on the occipital cortex, showed that the DAP12-deficient brain is not silent: it revealed a dysregulated microglial signature alongside reactive astrocytes and impaired-myelination oligodendrocytes^22^. What the NHD microglial state is, and through which secreted signals it reaches astrocytes, oligodendrocytes, and neurons, therefore remain open; and because NHD is white-matter-predominant and clinically frontal, a fixed microglial lesion might not produce the same state in brain regions of different anatomical and cellular composition.

To answer these questions, we profiled the prefrontal cortex and hippocampus from a *TYROBP* start-loss homozygote and a control donor by single-nucleus RNA sequencing, and the prefrontal cortex from both by spatial transcriptomics; this is the first study to combine the two in the NHD brain. We report four main findings. First, we define the NHD microglial state: DAP12-deficient microglia adopt a CIITA-driven, MHC class II-high antigen-presenting state whose upregulated receptors are Fc receptors rather than DAP12-coupled ones. In both regions, the TREM2-independent first step of the DAM trajectory is intact, and the TREM2-dependent second step is absent, which shows that the two steps described in mice are also separable in the human brain. Second, the mutant *TYROBP* transcript is retained in NHD microglia, so the lesion acts at the level of translation and the donor is DAP12-deficient rather than null. Third, from this state, the microglia remodel their secreted output onto astrocytes, which become pan-reactive and lose their metallothionein module, onto oligodendrocytes, whose maturation and cholesterol synthesis are impaired, and onto excitatory neurons, which lose presynaptic and activity-dependent programs; each change holds between the two regions and, for the glia, across the three NHD donors profiled to date. Fourth, in tissue, the microglial program concentrates in the layer I/pia and white-matter compartments, and microglia adopt the same state in both regions, while the response of their neighbors differs: broader in hippocampal astrocytes, greatest in prefrontal oligodendrocytes.

## Results

To determine how DAP12-deficient microglia influence the cells around them, and whether they do so alike in prefrontal cortex and hippocampus, we profiled both regions from one Nasu-Hakola disease (NHD) donor, a *TYROBP* c.2T>C (p.Met1Thr) start-loss homozygote (donor 861), and one control donor (CON) with no neurological diagnosis by single-nucleus RNA sequencing (snRNA-seq), and prefrontal cortex from both by spatial transcriptomics. Because the primary dataset is a single case-control pair, a consequence of a disease with approximately 200 reported cases worldwide, no claim rests on single-donor significance: per-nucleus tests only nominate genes and programs, and inference rests on effect size and on agreement in direction between the two regions and with the Zhou et al. cohort^22^ (3 NHD versus 11 CON, occipital cortex), judged against the empirical rate of chance agreement (Methods)^25^.

### Case presentation

The NHD donor, a Japanese woman homozygous for *TYROBP* c.2T>C (p.Met1Thr), followed the classical course of the disease (Table 1). The donor developed bone disease in her fourth decade, with cystic lesions and recurrent long-bone fractures; frontal dementia with disinhibition and loss of self-care began at age 40, and epilepsy at age 47. She died of pneumonia at age 59, after a 29-year course, with a brain weight of 700 g and the sclerosing leukoencephalopathy of the disease, together with intermediate Alzheimer-type change (Braak III, Thal 2). The sequence of bone, frontal and seizure manifestations is typical of NHD; the duration is longer than in most reported patients, so the tissue is end-stage. The control donor, a Japanese woman aged 55 with no neurological diagnosis, was obtained from the same brain bank.

**Table 1.**
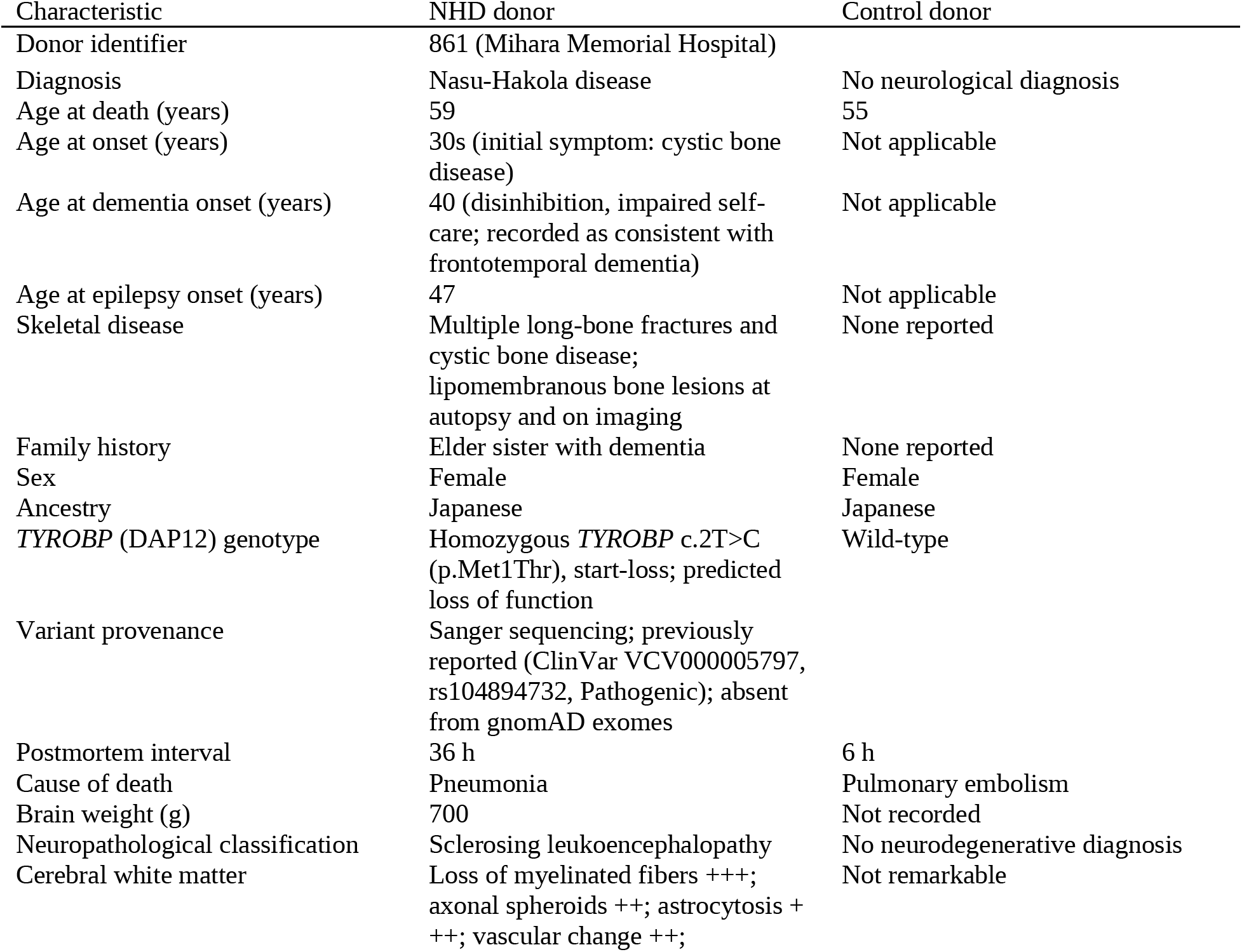
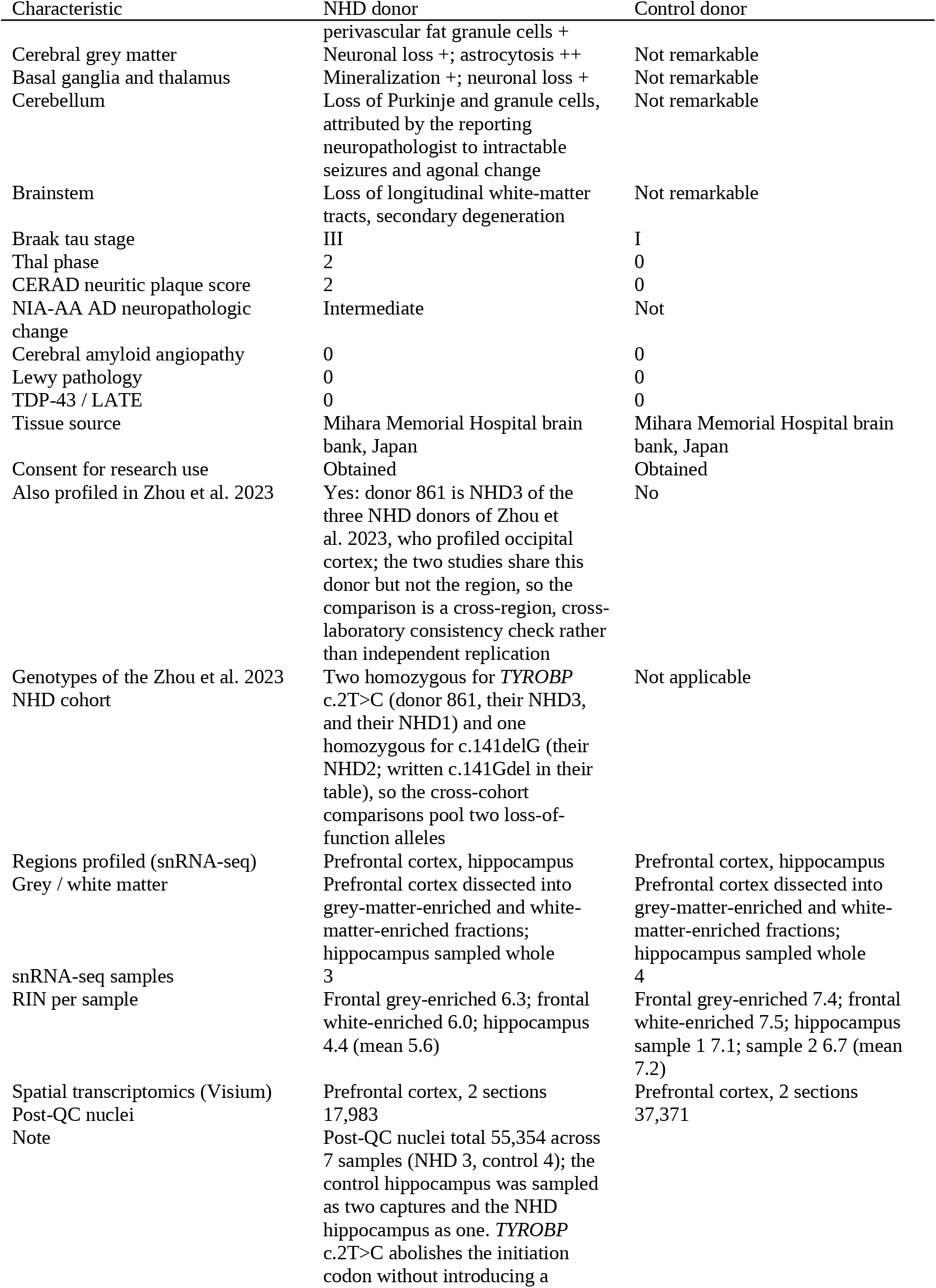
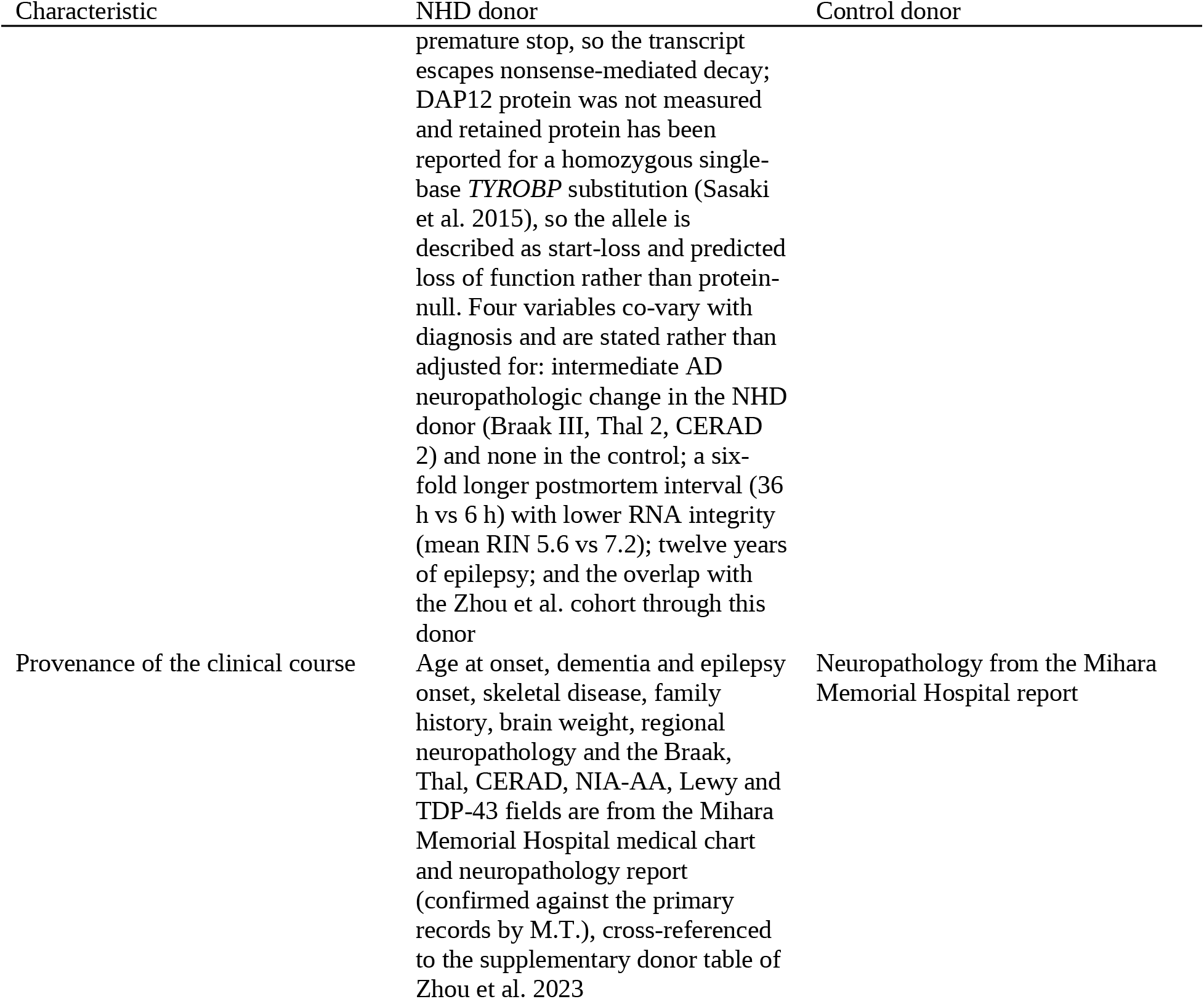
Donor and sample characteristics. The table lists diagnosis, age, sex, ancestry, *TYROBP* genotype and variant provenance, tissue source, regions and libraries profiled, Visium sections, and post-quality-control nuclei. Includes postmortem interval, cause of death, brain weight, the NHD donor’s clinical course (age at onset, bone disease, dementia, epilepsy, family history) and graded NHD neuropathology by region, Braak, Thal and CERAD scores, NIA-AA change, RIN per library, the donor’s inclusion in the Zhou et al. cohort with that cohort’s three NHD genotypes, and the provenance of each clinical field (Mihara Memorial Hospital records and neuropathology report, cross-referenced to the supplementary donor table of Zhou et al. 2023, which profiled occipital cortex from the same donor).

### The NHD response is partitioned by cell type and holds its direction between regions

To map the NHD response across cell types and brain regions, we profiled the prefrontal cortex and hippocampus, which were dissected, captured, and sequenced independently, from the NHD and the control donor. We recovered 55,354 post-quality-control nuclei from seven samples, four control and three NHD, spanning the prefrontal cortex, dissected into grey-matter-enriched and white-matter-enriched fractions, and the hippocampus, where the NHD tissue yielded a single sample (Fig. 1a; Supplementary Fig. 1; Supplementary Data 2, sheet Nucleus counts). Integration resolved eight canonical classes, microglia and perivascular macrophages (Micro-PVM), astrocytes, oligodendrocytes, oligodendrocyte precursor cells (OPC), excitatory and inhibitory neurons, endothelial cells and pericytes, that intermixed across condition, region, and sample (Fig. 1b; Supplementary Fig. 1b to f; Supplementary Fig. 5), with the NHD-enriched oligodendrocyte states described below (Fig. 4) as the one condition-restricted structure. The same eight classes were resolved in the Zhou cohort, which we used throughout to test cross-cohort direction consistency (Supplementary Fig. 2). With the cell types defined, we asked two questions of this dataset: how the NHD response is distributed across cell types and whether it is consistent between the two regions.

**Fig. 1.**
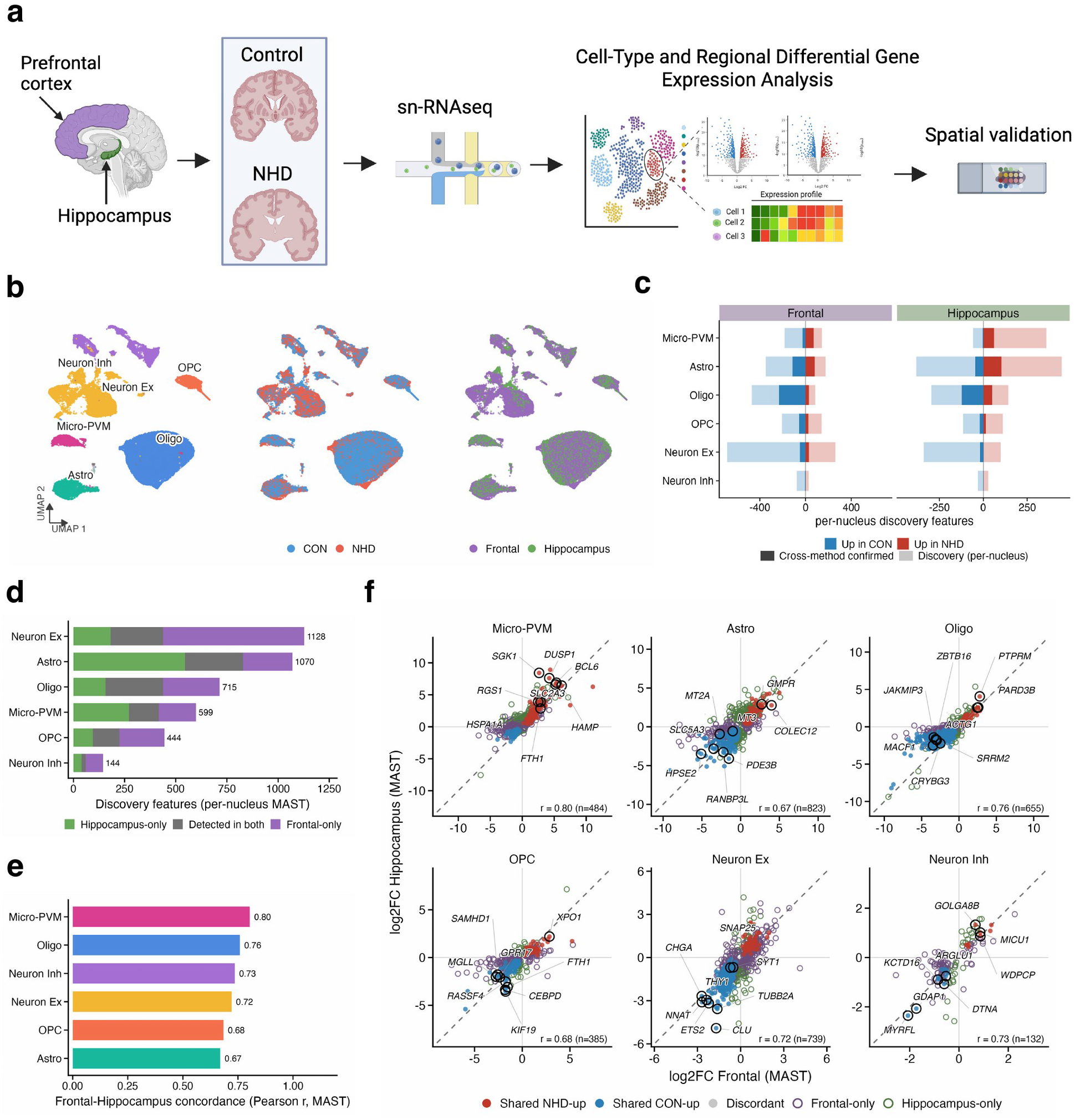
The NHD response is partitioned by cell type and direction-consistent between regions. **a** Experimental design. Prefrontal cortex, dissected into grey-matter-enriched and white-matter-enriched fractions, and whole hippocampus from one NHD and one control (CON) donor were profiled by single-nucleus RNA sequencing (snRNA-seq); prefrontal cortex from both donors was also profiled by spatial transcriptomics. Created in BioRender. Condello, C. (2027) https://BioRender.com/sszqbjh **b** UMAP embedding of the 55,240 nuclei carrying a canonical label, colored by cell type (left), condition (middle) and region (right). Endothelial cells and pericytes are plotted but not labeled. **c** Number of per-nucleus MAST discovery features per cell type and region (pale bars; Bonferroni-adjusted P < 0.05, |Cliff’s delta| ≥ 0.15, detection ≥ 10%), up in NHD (red) or in CON (blue), and the subset confirmed by frontal pseudobulk DESeq2 (solid bars). In hippocampus, which has no pseudobulk tier, a solid bar counts features whose sign agrees with the frontal pseudobulk sign (Methods). x scales are independent per region. **d** Discovery features per cell type partitioned into frontal-only, detected in both regions (any direction) and hippocampus-only; the union is printed at the end of each bar. **e** Pearson correlation of MAST log2 fold change between prefrontal cortex and hippocampus, computed per cell type over the genes tested in both regions (n per cell type in f). **f** The same correlations gene by gene: log2 fold change in hippocampus against prefrontal cortex. Filled discs, genes at discovery in both regions (shared NHD-up, shared CON-up or discordant); open purple and green rings, genes at discovery in one region only; black rings, the eight strongest genes per cell type, named; dashed line, identity. r and n are printed per panel. Correlations carry no significance test. Source data: Supplementary Data 2, 3, and 4; Table 1.

We tested differential expression in two steps: a discovery step at single-nucleus resolution, in which a per-nucleus test with an effect-size and detection gate nominated genes, and a confirmation step in which the nuclei of each sample were aggregated and the grey-matter-enriched and white-matter-enriched samples of each donor were treated as separate units (pseudobulk; Methods). We found that every cell type showed a transcriptional burden in NHD, and that its size differed by cell type more than by region (Fig. 1c). Excitatory neurons showed the most discovery features (946 in prefrontal cortex, of which 78 were confirmed at the pseudobulk level; 438 in hippocampus, 19 supported by the prefrontal direction), followed by astrocytes (524, 195 confirmed; 829, 149 supported), oligodendrocytes (558, 261; 438, 173), microglia (328, 97; 417, 65) and OPC (349, 80; 225, 34), with inhibitory neurons the least changed (105, 6; 59, 2). Pseudobulk confirmation was highest in the glia: 47% of prefrontal oligodendrocyte features and 37% of astrocyte features, compared with 8% of excitatory-neuron features, so the glial responses are the ones that hold at the level of the sample as well as the nucleus.

We next asked how each cell type’s response is distributed between the two regions, by splitting its features into those shared by the two regions and those found in one (Fig. 1d). Oligodendrocytes and astrocytes changed most regardless of region: they contained the largest shared cores (281 and 283 features, 39% and 26% of their unions), with near-complete direction agreement across the shared genes (277 of 281 and 263 of 283), such as the loss of *CALN1* and *TAFA5* in oligodendrocytes and of the metallothioneins *MT1G* and *MT1F* in astrocytes in both regions. Microglia shared 146 features (24% of 599; Supplementary Fig. 3a), including the iron regulator *HAMP* and the zinc transporter *TMEM163* up and the mannose receptor *MRC1* (CD206) down in both regions. Excitatory neurons, by contrast, changed most in a region-specific way: their union was the largest (1,128), but 690 features were prefrontal-only and only 256 shared, among them the loss of *NNAT*, *CHGA,* and *THY1*. The largest region-specific set belonged to hippocampal astrocytes (546 features in the hippocampus only), the one cell type with a greater burden in the hippocampus than in the prefrontal cortex.

Most features were discovered in only one region. Region-restricted detection can reflect region-specific biology or a single response in which different genes clear the gate in each region. To distinguish the two, we correlated the log2 fold changes of the two regions across the genes tested in both regions. We counted the discovery features whose signs agreed between regions, against the agreement expected by chance. We found that direction agreed in every cell type: Pearson r 0.67 to 0.80 over the discovery features, highest in microglia and oligodendrocytes (Fig. 1e), and sign agreement 93% to 100% against baselines of 67% to 84% (Fig. 1f), so region-restricted detection does not indicate region-restricted biology. The NHD response is therefore cell-type-partitioned in size, region-shared in the glia, and region-specific in the detected features for the neurons, but consistent in direction wherever the two regions overlap; this property frames the cell-type analyses that follow.

### DAP12-deficient microglia adopt a CIITA-driven antigen-presenting state that does not engage the TREM2-dependent arm

TREM2-DAP12 signaling is the switch that converts microglia from surveillance to a reactive state, so we asked what NHD microglia become when the switch is absent. Microglia were defined within the myeloid subset, from which re-clustering set aside a small CD163^+^/F13A1^+^ population of unresolved origin and T lymphocytes (Fig. 2a; Supplementary Fig. 3d to f; Supplementary Data 2, sheet Myeloid selection); the program scores and regulon activities below are computed on these 2,824 microglia as one population, the per-gene tests on the pooled myeloid class (Methods). We found that NHD microglia were nonetheless activated. In both regions, they adopted the same coherent state, a CIITA-driven, MHC class II-high antigen-presenting program, and did not progress into the TREM2-dependent arm of the DAM trajectory (Fig. 2b, d, h). At the level of single genes (Fig. 2b), NHD microglia downregulated the markers of the surveillant, homeostatic state (*P2RY12*, *MEF2C*) and upregulated, in both regions, genes of antigen presentation (*HLA-DRA*, *CD74*), complement production (*C1QB*), iron and zinc handling (*FTH1*, *TMEM163*), and glycolysis (*SLC2A3*). DAP12 is the adaptor through which TREM2 and its partner receptors signal, so the receptors that NHD microglia gained show how the state is sustained without it. We found that the upregulated receptors do not depend on DAP12 (Fig. 2c). Of the activating receptors that DAP12 couples to the kinase SYK, only *TREM2* was detected in human microglia (Supplementary Data 3). The most strongly upregulated receptors were the Fc receptors: depth-adjusted detection of *FCGR3A* and *FCGR1A*, which signal through the alternative ITAM adaptor FcRγ, and of *FCGR2A*, which carries its own ITAM, and *FCGR2B*, the inhibitory ITIM receptor, increased by 6 to 34 percentage points in the two regions, together with complement and scavenger receptors; the adaptors themselves changed little (*FCER1G* +3.7 and +1.2 points, *HCST* +1.8 and +1.3), so the gain is on the receptors, not the adaptors (Fig. 2c; Supplementary Data 12, sheet Compensatory routes). The transcript of the mutated adaptor itself was not lost but increased: *TYROBP* was detected in 18% and 25% of NHD microglia against 10% of control microglia in the two regions (log2 fold change +0.9 in both, below the discovery gate; depth-adjusted detection +6.5 and +0.5 points; Fig. 2c; Supplementary Data 3). A start-loss allele keeps its reading frame and introduces no premature termination codon. The retained message shows that the c.2T>C transcript escapes nonsense-mediated decay: the lesion acts at translation, not on the message. The microglia themselves were not depleted in the cortex: their spatial abundance was higher in every cortical layer of NHD prefrontal cortex (log2 NHD/CON +0.38 to +0.81) but not in the white matter, where it was lower (−0.54) alongside the loss of oligodendrocytes and their precursors (Fig. 6g; the nuclear fraction is not used for abundance, Methods).

**Fig. 2.**
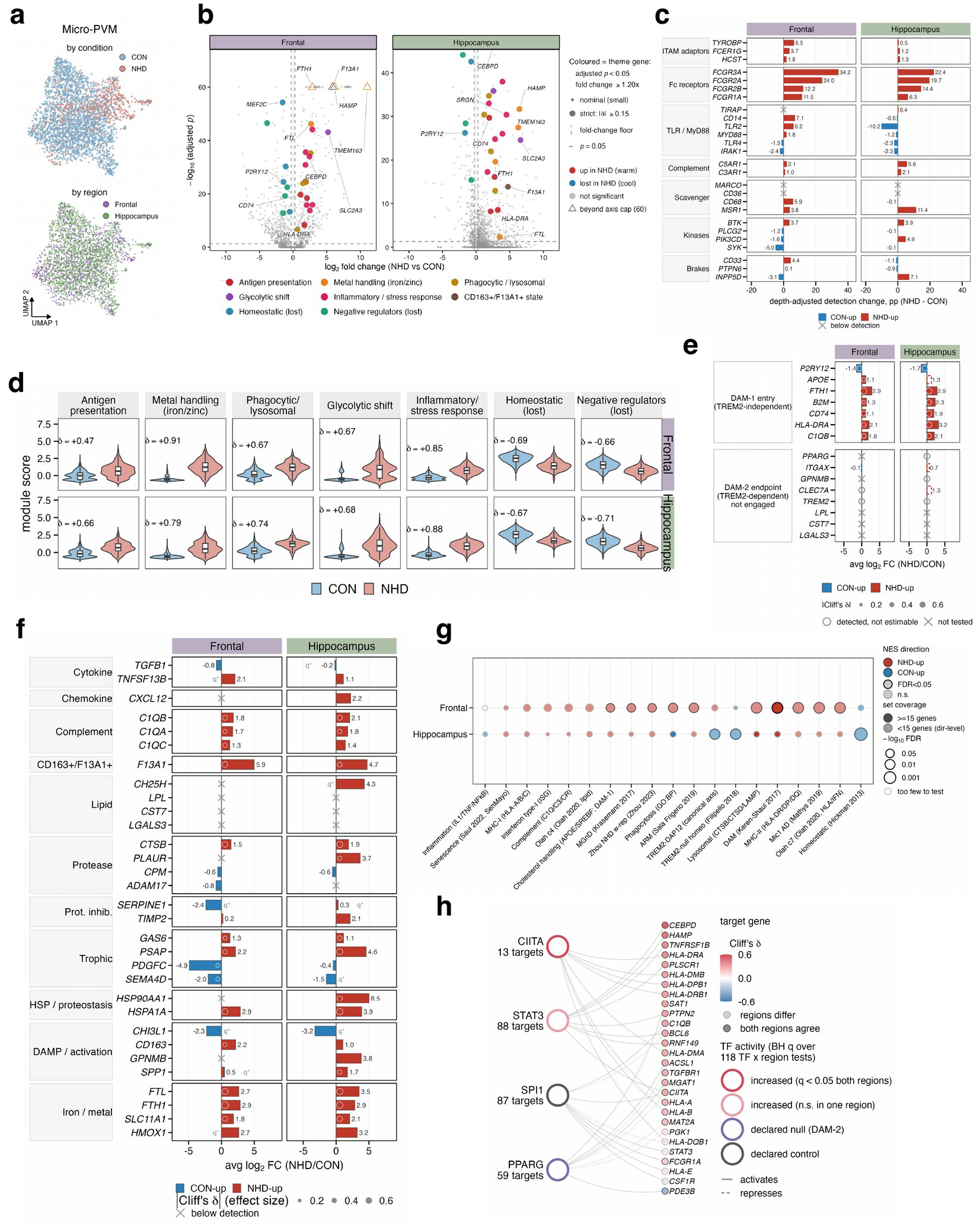
DAP12-deficient microglia adopt a CIITA-driven antigen-presenting state that does not engage the TREM2-dependent arm. **a** UMAP embedding of the microglial subset (2,824 nuclei after removal of the CD163^+^/F13A1^+^ population and T lymphocytes; Supplementary Data 2, sheet Myeloid selection), colored by condition (top) and region (bottom). **b** Volcano plots of per-nucleus MAST differential expression, NHD versus CON, per region. Colored points, genes of the named programs with Bonferroni-adjusted P < 0.05 and fold change ≥ 1.2; large points, |Cliff’s delta| ≥ 0.15; triangles, beyond the axis cap; dashed lines, fold-change floor and P = 0.05. **c** Depth-adjusted change in detection (percentage points, NHD minus CON) for receptor, adaptor, kinase and inhibitory-receptor genes, by family, on 2,986 myeloid nuclei; crosses, below the detection floor. **d** Violin plots of the seven microglial program scores per nucleus (microglia only), by condition and region; box, median and interquartile range; Cliff’s delta (NHD minus CON) printed per panel. **e** Log2 fold change of the disease-associated microglia (DAM) trajectory genes split into TREM2-independent entry genes and TREM2-dependent endpoint genes, per region; dot size, |Cliff’s delta|; cross, not returned by the test; open circle, detected but not reliably estimable (Supplementary Table S1). **f** Log2 fold change of secreted and membrane factors grouped by family, per region; dot size, |Cliff’s delta|; cross, below the detection floor. **g** Enrichment of NHD-versus-CON log2 fold change against 19 published microglial-state signatures, per region; fill, normalized enrichment score; dark ring, FDR < 0.05; faded, fewer than 15 set members detected in that region. **h** Regulon activity (decoupleR on the CollecTRI network) on 2,985 myeloid nuclei, Benjamini-Hochberg over 118 factor-by-region tests: CIITA and STAT3 pass the pre-declared gate, HIF1A is positive without passing, PPARG is a declared null and SPI1 an identity control (Methods). Target genes colored by Cliff’s delta; solid edge, activation; dashed edge, repression. Panels b, e, f and g use the pooled Micro-PVM class (3,081 nuclei); panels d and h the microglia-only set. Grey q*, Bonferroni-adjusted P < 0.05 with |Cliff’s delta| < 0.15. Source data: Supplementary Data 2, 3, 5, 6, 7, 8, and 12.

To read the gene-level changes as programs rather than as single genes, we scored each microglial nucleus for seven curated programs (Methods; Supplementary Data 6, sheet Signatures): antigen presentation, metal handling, phagocytic and lysosomal, glycolytic shift, inflammatory and stress response, the homeostatic program, and the negative regulators of activation (*IRAK3*, *LDLRAD4*, *ZBTB16*, *HDAC9*). We found that all seven changed in the same direction in both regions: the first five increased (Cliff’s delta +0.47 to +0.91) and the last two decreased (−0.66 to −0.71) (Fig. 2d; Supplementary Data 7, sheet Program deltas). The largest gains were in metal handling (+0.91 and +0.79) and in the inflammatory and stress response (+0.85 and +0.88). Metal handling was carried by the iron regulator *HAMP*, detected in 1% of control and 45% of NHD microglia, and the zinc transporter *TMEM163*, detected in 0% and 1% of control and in 61% and 52% of NHD microglia in the two regions. The glycolytic shift (+0.67 and +0.68) was confined to microglia (Methods). We then tested the prediction of the two-step model directly, by splitting the DAM trajectory into its TREM2-independent entry genes and its TREM2-dependent endpoint genes (Fig. 2e). The two arms separated: every TREM2-independent entry gene changed in the expected direction, all but hippocampal *APOE* at the discovery level, whereas no TREM2-dependent DAM-2 gene reached discovery in either region (Supplementary Table S1). In the prefrontal cortex the DAM-2 effect sizes were null (*CLEC7A* Cliff’s delta +0.09, *ITGAX* −0.02), and *LPL*, *CST7*, *LGALS3*, *GPNMB* and *PPARG* were detected in fewer than 10% of microglia in both conditions, below the floor at which the test is run, so they could not be scored (crosses in Fig. 2e). In the hippocampus, a single NHD library, *CLEC7A* and *ITGAX* trended upward (Cliff’s delta +0.21 and +0.15) without reaching significance, the same status as hippocampal *APOE*.

This state also changed what the microglia secrete (Fig. 2f): complement (*C1QA/B/C*), iron and metal factors (*FTL*, *FTH1*, *SLC11A1*), the trophic ligand *GAS6* and the TNF-family cytokine *TNFSF13B* (BAFF; +2.1 and +1.1 log2 fold change) were upregulated, together with the TNF receptor *TNFRSF1B* (discovery in both regions; Supplementary Data 3), whereas *TGFB1* was slightly downregulated; *SPP1* was upregulated in hippocampus and, on microglia alone, in prefrontal cortex (Supplementary Data 8, sheet Pooled vs microglia-only). To place this state among published ones, we scored enrichment against nineteen microglial-state signatures (Fig. 2g): the NHD prefrontal microglia resembled the DAM, MHC-II-high, activated-response and MGnD states, and in hippocampus the homeostatic-identity and TREM2-DAP12-axis sets were enriched on the control side; the one lipid signature enriched on the NHD side (lipid/cholesterol, false discovery rate, FDR, 0.04) is carried by *APOE* and the TREM2-independent entry genes, not by the TREM2-dependent set. Genome-wide pathway enrichment returned the same complement-, MHC- and Fc-receptor-carrying immune terms: lymphocyte-mediated immunity (normalized enrichment score, NES, 2.19; FDR 0.006), a term carried here by *FCGR3A*, *HLA-DRA*, *C1QA/B/C*, *B2M*, *FCGR1A*, *HLA-DRB1*, *CD74* and *TYROBP* rather than by lymphocyte genes, antigen processing and presentation (2.03, 0.047) and regulation of the inflammatory response (1.86, 0.047), with no term lost (Supplementary Fig. 3b). Finally, to identify the transcription factors behind this state, we estimated regulon activity for 59 factors in each region against a pre-declared gate (Methods; Fig. 2h), with SPI1 (PU.1), the lineage-determining transcription factor of microglia, as identity control: its regulon should not move if the population keeps its identity, and it did not (q = 0.34 and 0.72). The MHC class II transactivator CIITA passed the gate in both regions (Benjamini-Hochberg q = 0.041 and 8.4 × 10^−5^), as did STAT3, whose activity was positive in both regions and significant in prefrontal cortex (q = 0.006 and 0.051). HIF1A, the transcriptional driver of glycolysis, was positive in both regions without passing the gate (q = 0.24 and 0.071), so the glycolytic shift is read from its genes rather than from its regulator. PPARG, the transcription factor of the TREM2-dependent lipid arm, showed no reproducible activity: +0.70 in prefrontal cortex (q = 0.78) and +0.02 in hippocampus (Fig. 2h).

The NHD microglial program agreed in direction with the Zhou et al. cohort, and every cell type in both regions exceeded its empirical rate of chance agreement, beyond chance in ten of the twelve comparisons (Supplementary Data 9, sheet Direction consistency): 75% of 72 microglial genes significant in both datasets agreed in prefrontal cortex and 82% of 67 in hippocampus, against baselines of 50.0% and 50.9% (binomial P = 1 × 10^−5^ and 1 × 10^−7^; Supplementary Fig. 3c). Together these data define the NHD microglial state as a CIITA-driven, antigen-presenting program whose upregulated receptors are Fc receptors rather than DAP12-coupled ones, with the TREM2-dependent arm of the DAM trajectory^19^ not engaged in either region.

### NHD microglia rewire secreted signaling onto astrocytes and the oligodendrocyte lineage

A changed secretome in NHD microglial cells (Fig. 2f) could further influence neighboring glial cells. We therefore tested whether the NHD microglial output could reach astrocytes and the oligodendrocyte lineage. We predicted ligand-receptor interactions with CellChat, which scores each pair by the co-expression of the ligand in the sender and its receptor in the receiver as a communication probability and sums the pairs of a family into a pathway (Methods). We found that NHD microglia increased their outgoing signaling onto both receivers. Onto astrocytes, the largest gain was the SPP1 pathway, whose probability increased from 0.039 to 0.048 in prefrontal cortex and from 0.041 to 0.145 in hippocampus; at pair level the gain was carried by SPP1-CD44 (change in communication probability +0.028 and +0.098) and GAS6-MERTK (+0.026 and +0.005) (Fig. 3g; Supplementary Data 12, sheet LR microglia to glia). The GAS6 gain met a downregulated receptor: *MERTK*, the TAM receptor for GAS6 on astrocytes, was downregulated at the effect-size gate in hippocampus (Cliff’s delta −0.30). Onto the oligodendrocyte lineage, microglial input lost the *TENM4*-*ADGRL3* pair in prefrontal cortex (−0.113 on OPC), gained *PDGFB*-*PDGFRA* in hippocampus (+0.036) and kept *PSAP*-*GPR37* onto oligodendrocytes in both regions (Fig. 4j; Supplementary Data 12, sheet LR microglia to glia).

**Fig. 3.**
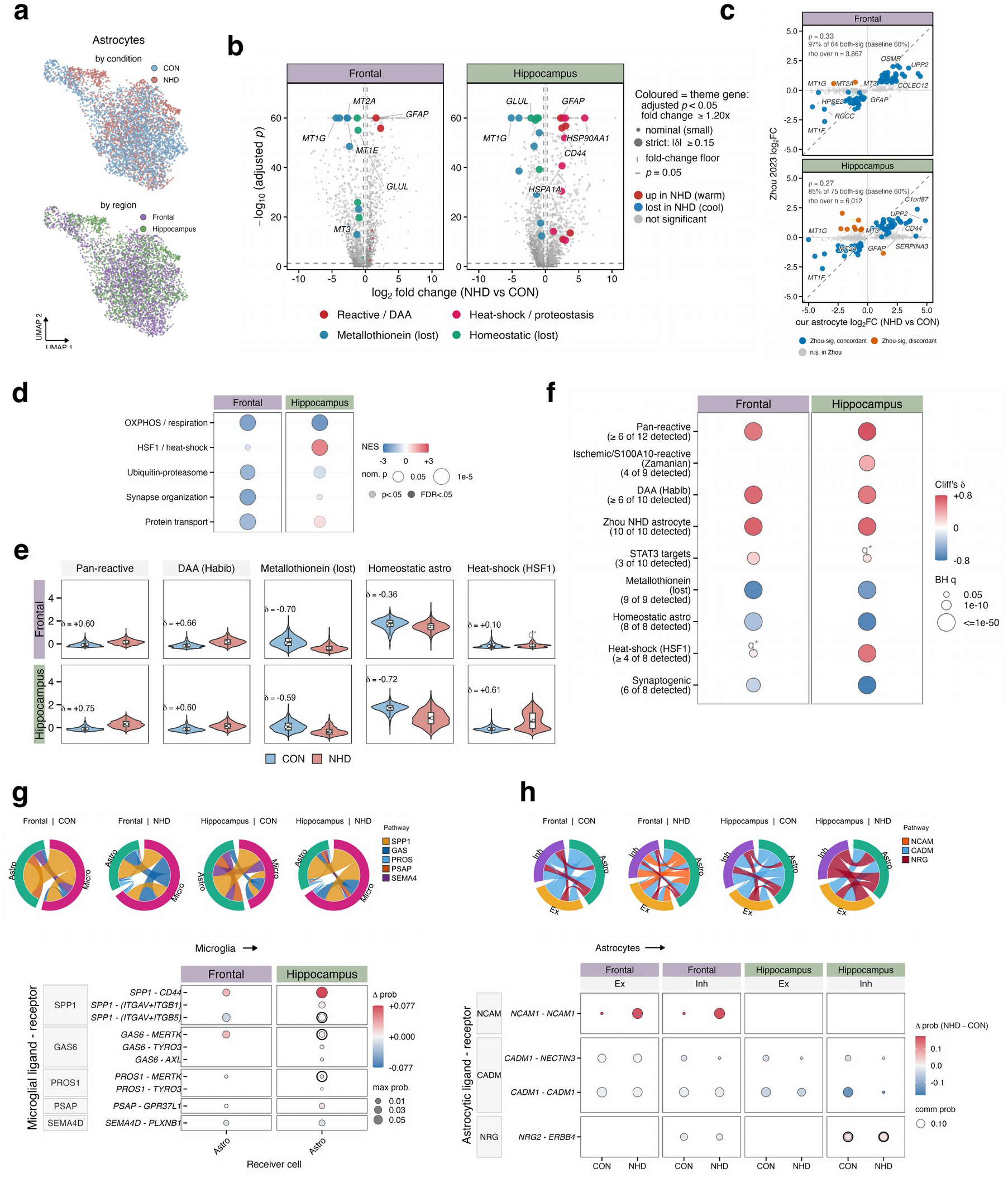
Astrocytes adopt a pan-reactive program and lose the metallothionein module. **a** UMAP embedding of the astrocyte subset (4,978 nuclei), colored by condition and region. **b** Volcano plots of per-nucleus MAST differential expression per region, as in Fig. 2b; colored points, genes of the reactive, metallothionein, heat-shock, and homeostatic programs. **c** Agreement in direction with the Zhou et al. cohort (occipital cortex; not region-matched): our astrocyte log2 fold change (x) against theirs (y), per region. Blue, same direction in both cohorts; vermilion, opposite; grey, not significant in Zhou et al. The fraction of both-significant genes agreeing in direction, its empirical baseline across all shared genes, and the Spearman rho are printed per panel; a one-sided binomial test against the baseline (Methods). **d** Gene-set enrichment per region; fill, normalized enrichment score; dot size, nominal P; dark ring, FDR < 0.05. **e** Violin plots of five signature scores per nucleus, by condition and region; box, median and interquartile range; Cliff’s delta printed per panel. **f** Cliff’s delta (NHD minus CON) for the astrocyte signatures per region (eleven tested in each region; the nine shown passed the detection gate in at least one region); fill, delta; dot size, Benjamini-Hochberg q; row labels give the number of set members detected in 10% of nuclei, and a cell below that gate is not drawn. Signatures are scores of one population, not subtypes. **g** Microglia-to-astrocyte communication (CellChat). Top, chord diagrams of pathway-level communication per condition and region; bottom, change in communication probability (NHD minus CON) per ligand-receptor pair; dot size, maximum probability. **h** Astrocyte-to-neuron communication for the four pairs whose astrocytic ligand clears the ambient-expression filter applied to the volcanoes, per condition, region and receiver class; fill, change in probability; dot size, probability. In g and h a black ring marks a receptor lost in the receiver at discovery level, and a blank cell is a pair not inferred in that region; communication probabilities are effect sizes only. Grey q*, per-nucleus Wilcoxon BH q < 0.05 within panel (22 tests) with |delta| < 0.15; NHD hippocampus is one sample. Source data: Supplementary Data 3, 5, 6, 7, 9, and 12.

**Fig. 4.**
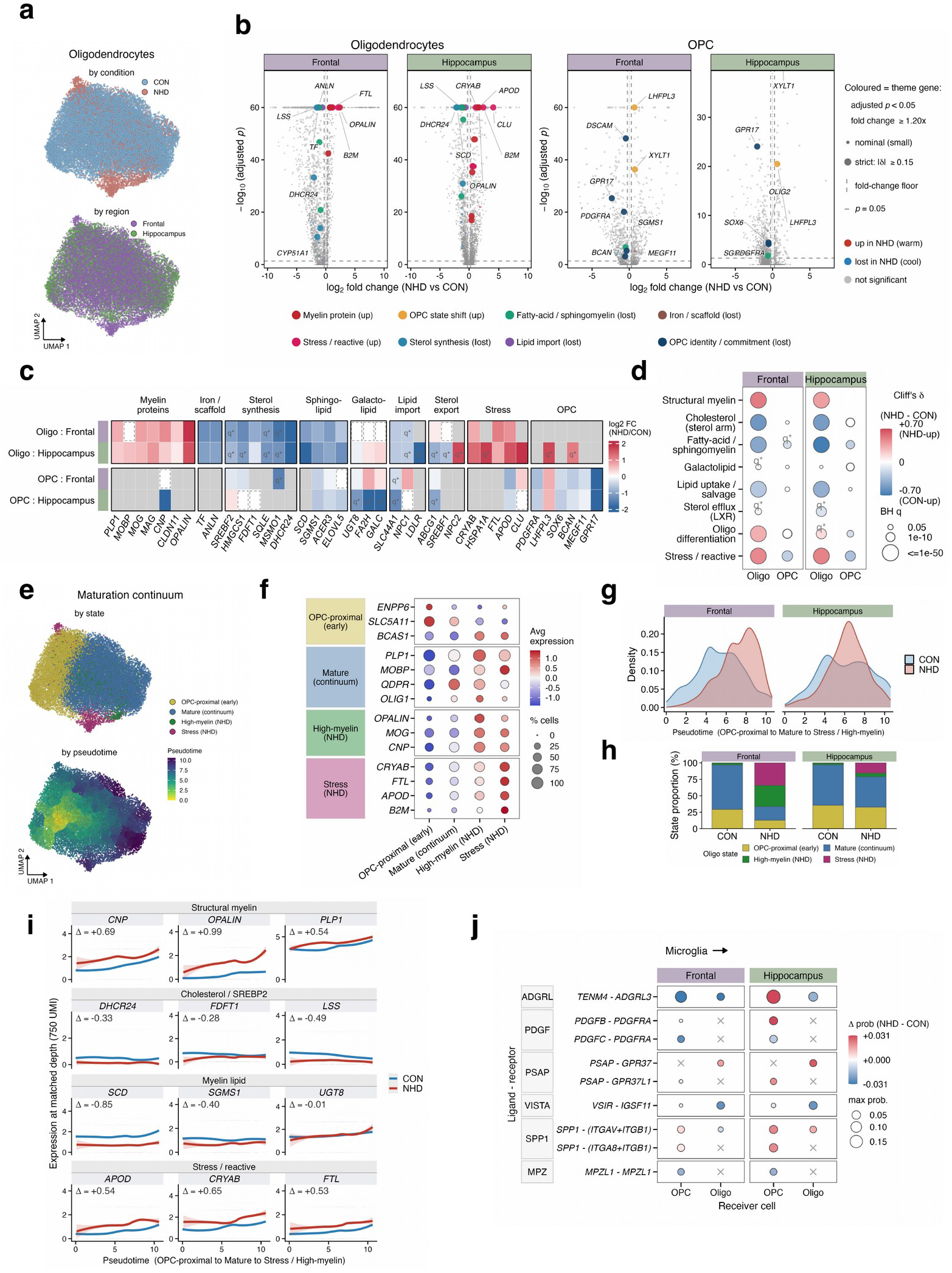
Oligodendrocyte maturation and cholesterol synthesis are impaired in NHD. **a** UMAP embedding of the oligodendrocyte subclustering (25,317 nuclei), colored by condition and region; OPC were analyzed separately. **b** Volcano plots of per-nucleus MAST differential expression for oligodendrocytes and OPC per region, as in Fig. 2b; colored points, genes of the myelin, stress, lipid-synthesis, lipid-import, iron/scaffold and OPC programs. **c** Heat map of log2 fold change per gene, arranged by pathway block, in four lineage-by-region tracks; grey, below the 10% detection floor; dashed outline at zero, detected but stable; q*, Bonferroni-adjusted P < 0.05 with |delta| < 0.15 (Supplementary Data 3). **d** Cliff’s delta (NHD minus CON) for eight lineage programs in oligodendrocytes and OPC per region; fill, delta; dot size, Benjamini-Hochberg q. A cell is drawn only where three set members, or two covering half the set, reach 10% detection; hollow, clears the effect gate on share but not at matched depth. **e** UMAP of the maturation continuum colored by state (top) and by Monocle 3 pseudotime (bottom; yellow early to purple late). **f** Dot plot of state markers; dot size, percent of nuclei expressing; fill, scaled mean expression. **g** Density of nuclei along pseudotime by condition and region. **h** State composition per condition and region (frontal CON 14,588 and NHD 2,303 nuclei; hippocampus 7,081 and 1,345). The two NHD-enriched states were defined in part by NHD enrichment (Methods); the depletion of mature oligodendrocytes was not. **i** Expression along pseudotime at matched depth (750 UMI per nucleus; 22,411 nuclei) by condition; line, smoothed mean; band, dispersion across nuclei; delta, mean NHD-minus-CON difference along the trajectory. **j** Microglial input onto OPC and oligodendrocytes (CellChat): change in communication probability per pair and region; dot size, maximum probability; cross, receptor not detected; color scale clipped at ±0.031. Grey q*, per-nucleus Wilcoxon BH q < 0.05 within panel (32 tests) with |delta| < 0.15; NHD hippocampus is one sample. Source data: Supplementary Data 3, 5, 6, 7, and 12.

### Astrocytes adopt a pan-reactive program with loss of the metallothionein module

Astrocytes can change their state in response to microglial signals, and crosstalk with microglia shapes astrocyte reactivity across inflammatory and degenerative conditions of the brain^26^. Reactive astrocytes are no longer classified as simply neurotoxic or neuroprotective, but along several independent axes^27^. We scored each reactive axis in every astrocyte nucleus (4,978; Fig. 3a) and asked along which axes the NHD astrocytes changed. We found that NHD astrocytes became reactive along the pan-reactive and disease-associated axes in both regions, and at the same time lost the metallothionein module, a loss in the opposite direction to the rest of the reactive program. The per-gene changes fell into four groups (Fig. 3b): pan-reactive genes were upregulated in both regions (*GFAP* +1.61 and +2.30 log2 fold change); heat-shock genes were upregulated in hippocampus only (*HSP90AA1* +5.91); the metallothioneins were the most strongly downregulated (*MT2A* −2.70, Cliff’s delta −0.61, in prefrontal cortex; *MT1G*, *MT1E*, *MT1F* and *MT3* down in both regions); and the genes of homeostatic identity were downregulated (*GLUL* −1.10 and −2.16). These directions were shared with the Zhou et al. cohort: of the genes significant in both datasets, 97% of 64 agreed in prefrontal cortex and 85% of 75 in hippocampus, against baselines of 59.8% and 59.6% (binomial P = 4.7 × 10^−12^ and 1.2 × 10^−6^; Fig. 3c), and the agreement held in both regions when the shared donor was removed from their cohort (Supplementary Fig. 2e). Pathway enrichment placed oxidative phosphorylation below control in both regions (normalized enrichment score −2.01 and −2.29), ubiquitin-proteasome and synapse-organization terms below control in prefrontal cortex only, and the HSF1 heat-shock response above control in hippocampus only (+2.07, FDR 1.0 × 10^−4^; Fig. 3d). Scored as signatures, the pan-reactive (Cliff’s delta +0.60 and +0.75), disease-associated astrocyte^28^ (+0.66 and +0.60) and Zhou NHD astrocyte (+0.69 and +0.68) programs increased in both regions, and the metallothionein (−0.70 and −0.59), homeostatic (−0.36 and −0.72) and synaptogenic (−0.25 and −0.77) programs decreased in both, so the reactive gain and the metallothionein loss occur together in both regions (Fig. 3e, f). Within the metallothionein module, *MT3* merits separate mention: it is the growth-inhibitory factor first identified as deficient in the Alzheimer’s disease brain^29^, it buffers zinc and copper, and its loss is one of the features that separate the NHD astrocyte from a generic reactive astrocyte. Astrocyte-to-neuron signaling was scored for the four pairs whose astrocytic ligand clears the ambient-expression filter applied to the volcanoes (Methods). It was retained in prefrontal cortex, where no pair lost more than 0.03 in communication probability and *NCAM1*-*NCAM1* was gained from a zero baseline (+0.17 onto excitatory and +0.18 onto inhibitory neurons), whereas *CADM1*-*CADM1* decreased in hippocampus (−0.05 excitatory, −0.14 inhibitory) (Fig. 3h; Supplementary Data 12, sheet LR astrocyte to neuron).

### Oligodendrocyte maturation and cholesterol synthesis are impaired in NHD

White-matter degeneration is the defining brain lesion of NHD; the sclerosing leukoencephalopathy gives the disease its name and its radiological signature^5^, and this donor showed the lesion in its complete form (Table 1). A leukoencephalopathy can arise from a loss of mature oligodendrocytes, from a failure of the myelination program in the cells that remain, or from both. We therefore determined how the oligodendrocyte lineage is dysregulated in the presence of DAP12-deficient microglia. NHD and CON oligodendrocytes intermixed on subclustering (Fig. 4a), a graded shift rather than a segregated disease population; OPC were analyzed separately and carry a phenotype of their own. We found that NHD oligodendrocytes expressed more myelin-protein transcripts and fewer cholesterol-synthesis transcripts. At the level of single genes (Fig. 4b), two arms separated: myelin-protein and stress genes were upregulated (*OPALIN* +2.39 and +1.58 log2 fold change) while a coordinated lipid-synthesis arm was downregulated (*DHCR24* −2.05 and −2.19, *LSS*, *SCD*), and in OPC the receptor *PDGFRA* and *GPR17*, an inhibitor of oligodendrocyte maturation, were downregulated (−0.72 and −2.35 log2 fold change in prefrontal cortex). Arranging the tested genes by pathway block (Fig. 4c) showed the same split across the whole program: the myelin-protein block held flat to up, whereas the sterol, sphingolipid and lipid-import blocks were downregulated together with the galactolipid block flat, and the transcription factor of the sterol arm was downregulated with its targets: *SREBF2* −1.09 and −0.78 log2 fold change, its feedback sensor *INSIG1* −0.77 and −1.21, and *LDLR* downregulated in hippocampus (detected in 14% of control and 2% of NHD oligodendrocytes). The SREBP2 cholesterol-synthesis cascade was thus the most coherent single decline, the expected direction for a lesion of myelin membrane growth^30^. Iron handling was inverted in the same cells: the transferrin gene *TF was* downregulated (−1.31 and −1.09), while the ferritin heavy chain *FTH1* was upregulated (+1.14 and +1.15), and the scaffold gene *ANLN* was downregulated.

To score these arms per nucleus, we defined eight lineage programs, structural myelin, oligodendrocyte differentiation, the cholesterol, fatty-acid and sphingomyelin, and galactolipid synthesis arms, lipid uptake and sterol efflux, and stress (Methods; Supplementary Data 6, sheet Signatures), and scored them in oligodendrocytes and OPC of both regions (Fig. 4d). In oligodendrocytes, structural myelin increased (Cliff’s delta +0.51 and +0.38). In contrast, cholesterol synthesis (−0.54 and −0.51), fatty-acid and sphingomyelin synthesis (−0.45 and −0.68) and lipid uptake, carried by the choline transporter *SLC44A1* (−0.28 and −0.41), decreased; galactolipid synthesis and sterol efflux did not move (|delta| ≤ 0.14); and the stress program increased (+0.46 and +0.49). In OPC, the stress program changed in the opposite direction (−0.24 and −0.32), and no other OPC program cleared the effect gate at matched depth. Two interpretations of this dissociation are possible. The cells that remain may have changed their program, or they may be in a different state along the maturation continuum. To separate the two, we performed trajectory analysis. Each oligodendrocyte-lineage nucleus was placed along a maturation continuum by pseudotime, which orders nuclei by transcriptome from the OPC-like to the fully mature program, and states along the continuum were then assigned from their markers (Methods). We found that the continuum held four positions: OPC-proximal, mature, and two NHD-enriched states, a high-myelin (*OPALIN*^+^) state and a stress (*CRYAB*^+^) state, each with its own markers (Fig. 4e, f). The NHD nuclei were displaced from the mature center of the trajectory (Fig. 4g). The shift in composition was large in prefrontal cortex: mature oligodendrocytes decreased from 67.6% to 21.0% of the lineage and OPC-proximal cells from 29.2% to 12.7%, while the high-myelin and stress states increased from 2.9% to 32.0% and from 0.2% to 34.3% (Fig. 4h; Supplementary Data 12, sheet Oligo state composition); the hippocampal shifts ran the same way but smaller. We then asked whether the dissociation also holds within the cells that remain by comparing expression along pseudotime at matched sequencing depth (Fig. 4i). The deficit persisted: myelin genes remained above control levels across the trajectory, while the cholesterol and myelin-lipid genes remained below control levels. Together, these analyses show that NHD impairs oligodendrocyte maturation. Mature oligodendrocytes are depleted, the remaining cells occupy two NHD-enriched states, and the cholesterol-synthesis deficit persists within them; the per-cell increase in myelin transcript is a share of each surviving cell’s transcriptome, not an amount of tissue myelin, and the two quantities move in opposite directions because the denominator changes. The lesion is therefore compositional, a loss of mature oligodendrocytes with a coordinated cholesterol-synthesis deficit, rather than a failure of myelin transcription in the cells that remain, and we describe the lineage as one continuum plus two NHD states rather than as discrete disease subtypes ^31^. The oligodendrocyte program agreed in direction with the Zhou et al. cohort above its chance rate in both regions (65% of 31 genes against a 48% baseline in prefrontal cortex, binomial P = 0.046; 78% of 40 against 54% in hippocampus; Supplementary Data 9, sheet Direction consistency).

### Excitatory neurons downstream of the glial cascade lose presynaptic and activity-dependent programs, and oxidative phosphorylation in the prefrontal cortex

NHD ends in dementia, so the neuronal programs that are lost or become dysfunctional, and the glial input that could cause this dysfunction or impairment, are the last link in the cascade we aimed to elucidate. We examined neurons at three levels: gene-by-gene, as programs scored per nucleus, and as receivers of glial input. Neurons were separated into excitatory and inhibitory classes within the neuronal subset (17,963 nuclei; Fig. 5a), and each class was tested per region.

**Fig. 5.**
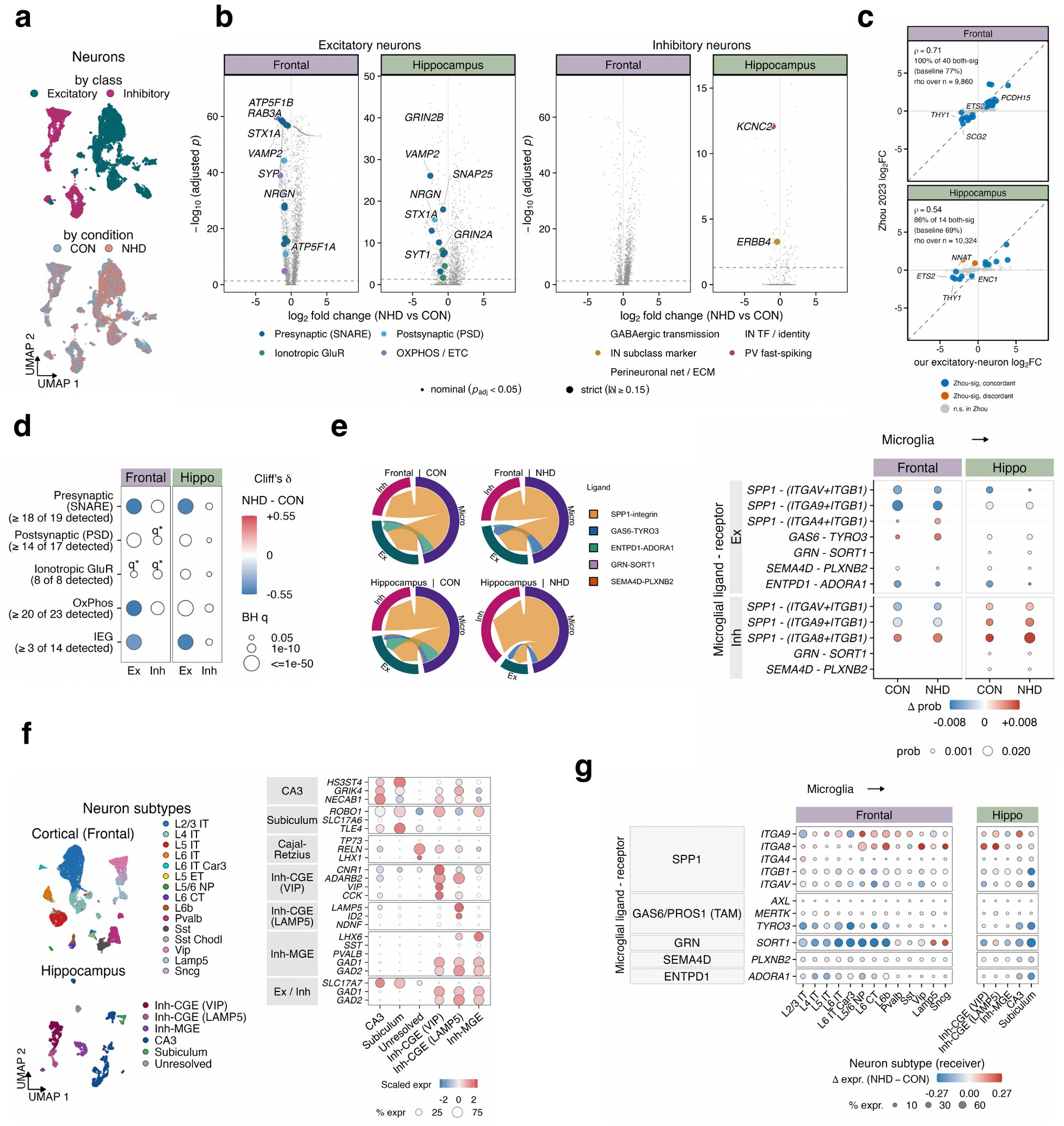
Excitatory neurons lose presynaptic and activity-dependent programs downstream of microglial input. **a** UMAP embedding of the neuronal subset (17,963 nuclei; CON 7,233, NHD 10,730), colored by class (top) and condition (bottom). **b** Volcano plots of per-nucleus MAST differential expression for excitatory and inhibitory neurons per region, as in Fig. 2b; colored points, genes of the synaptic, glutamate-receptor, oxidative-phosphorylation, and interneuron programs. No inhibitory program gene moves in the expected direction above the fold-change floor, so those facets carry no colored points. **c** Agreement in direction with the Zhou et al. cohort (occipital cortex; not region-matched) for excitatory neurons, as in Fig. 3c. **d** Cliff’s delta (NHD minus CON) for five neuronal programs in excitatory (Ex) and inhibitory (Inh) neurons per region; fill, delta; dot size, Benjamini-Hochberg q. Row labels give the number of set members detected in 10% of nuclei; a cell below that gate is not drawn; hollow, within the loss that sequencing depth alone can produce (Methods). **e** Microglia-to-neuron communication (CellChat). Left, chord diagrams per condition and region; right, change in communication probability per pair onto excitatory and inhibitory neurons; dot size, probability. **f** Neuronal taxonomy: UMAP of 15 cortical subclasses and five hippocampal subtypes plus one unresolved cluster (grey; 190 nuclei), with a dot plot of their marker genes; fill, scaled expression; dot size, percent expressing; hollow dots, detection below 10%. **g** Change in receptor expression (NHD minus CON, normalized mean) on each receiving subtype for the microglial ligands in e; dot size, percent of nuclei expressing; cells with fewer than 20 nuclei per condition dropped. Grey q*, per-nucleus Wilcoxon BH q < 0.05 within panel (20 tests) with |delta| < 0.15; NHD hippocampus is one sample. Source data: Supplementary Data 2, 3, 6, 7, 9, and 12.

We found that excitatory neurons lost presynaptic and activity-dependent programs in both regions, and oxidative phosphorylation in the prefrontal cortex, beyond what sequencing depth alone accounts for, and that the prefrontal cortex loss agreed in direction with the Zhou et al. cohort. At the level of single genes (Fig. 5b), the largest losses were synaptic and mitochondrial transcripts (*VAMP2*, *NRGN*, *SYT1*, *RAB3A* and *ATP5F1B* in prefrontal cortex; *GRIN2B*, *VAMP2* and *STX1A* in hippocampus), and under the discovery gate 74% of 946 prefrontal and 79% of 438 hippocampal excitatory features were downregulated in NHD. The prefrontal excitatory response agreed in direction with the Zhou et al. cohort, 100% of 40 genes significant in both datasets agreeing against a 77.3% baseline over 9,860 shared genes (P = 3.4 × 10^−5^). In contrast, the hippocampal arm (86% of 14 against 69.4%) did not exceed its baseline (P = 0.15) (Fig. 5c).

To read these losses as programs, we scored each neuron for synaptic transmission, oxidative phosphorylation, and immediate-early-gene activity, the energetically expensive functions that fail first across neurodegenerative diseases. Excitatory neurons lost the presynaptic program (Cliff’s delta −0.54 prefrontal, −0.51 hippocampus). Immediate-early genes (−0.42 and −0.52), each beyond the loss that sequencing depth alone produces (Methods) in both regions, and oxidative phosphorylation in prefrontal cortex (−0.55, against a loss of 0.29 from depth alone); in inhibitory neurons no program loss exceeded what depth alone produces (Fig. 5d; Supplementary Data 7, sheet Program deltas). Sterol biosynthesis was also depleted in excitatory neurons, significantly in the prefrontal cortex, with a concordant trend in the hippocampus (normalized enrichment score −1.59 and −1.53; FDR 0.04 and 0.06; Supplementary Data 5), so the cholesterol-synthesis deficit is not confined to the oligodendrocyte lineage.

The microglial secretome changed (Fig. 2f) and the neuronal programs decreased; to connect the two, we determined which microglial ligands reach neurons and which neurons receive them. We found that the microglial input onto neurons changed in two opposite directions rather than as a uniform gain (Fig. 5e): *GAS6*-*TYRO3* input increased onto prefrontal excitatory neurons (+0.0054) and *SPP1*-integrin input onto hippocampal inhibitory neurons (+0.0144), while the inhibitory purinergic pair *ENTPD1*(CD39)-*ADORA1* was lost (−0.0058; Supplementary Data 12, sheet LR microglia to neuron). To identify the receivers, we resolved the neuronal taxonomy into 15 cortical subclasses and five hippocampal subtypes (Fig. 5f; Methods). Among the cortical subclasses, the deep-layer excitatory neurons (L6 IT, L6 CT, L6 IT Car3 and L6b) were under-represented in NHD in both the grey- and the white-matter capture, whereas L4 IT held a similar share in both conditions (21.5% and 21.0% of excitatory nuclei; Supplementary Data 2, sheet Frontal subclasses); dentate granule neurons were not captured, and CA1 and CA2 did not separate from the CA3-marker cluster (Methods). At subtype resolution the receptors for the two upregulated microglial ligands were downregulated on the deep-layer receivers: the *SPP1*-integrin receptors *ITGAV* (−0.21) and *ITGB1* (−0.11) and the TAM receptor *TYRO3* (−0.26) were downregulated on prefrontal L6 corticothalamic neurons, with *AXL* undetected and *MERTK* flat (Fig. 5g; Supplementary Data 12, sheet LR microglia to neuron subtype), and at pair level the canonical SPP1-ITGAV/ITGB1 route decreased onto excitatory neurons in both regions while SPP1-ITGA8/ITGB1 increased onto inhibitory neurons. Astrocyte-to-neuron synapse-organizing input, by contrast, was broadly preserved in prefrontal cortex (Fig. 3h); of the glial inputs to neurons that we modeled, the changes were therefore microglial rather than astrocytic.

### The glial response is shared across the three NHD donors profiled to date

Because our donor is also NHD3 of the Zhou et al. cohort, agreement between the two studies could be the same brain measured twice. We therefore recomputed their NHD-versus-control contrast with the NHD side restricted by donor, all three donors, each leave-one-donor-out pair and each donor alone, against the same controls, and re-scored the agreement with our fixed effects in each arm; this separates what is shared across individuals from what varies between them (Supplementary Fig. 2e; Supplementary Data 9, sheet Donor arms). We found that the glial response was shared. With the two donors not shared with this study, microglial agreement in prefrontal cortex was unchanged (76% of 55 genes against a 51% baseline; Spearman rho 0.25 in both analyses) and astrocytic agreement persisted (92% of 53 genes; rho 0.36 to 0.16). The neuronal response varied between individuals and regions: without our donor the excitatory-neuron correlation decreased from 0.75 to 0.41, the two other donors disagreed with each other on neurons within the Zhou cohort itself (rho 0.17), one agreed with ours (0.57) and the other did not (0.01), and the comparison is in any case occipital cortex against prefrontal cortex and hippocampus. We therefore read the neuronal cross-cohort agreement as consistency within our donor across regions and laboratories, and the glial programs as the conserved core of the NHD response across the three unique NHD donors represented in the two studies: this donor in two regions and two laboratories, and the two independent donors of Zhou et al.

### Spatial transcriptomics localizes the glial cascade *in situ*

Single-nucleus data locate a program in a cell type but not in the tissue. We next used spatial transcriptomics to place the programs of the single-nucleus analysis in intact prefrontal cortex, to test whether they localize where the cascade predicts and whether they are graded by local microglial content. Across four Visium sections (two per donor; 18,765 spots), expression and neighborhood-aware clustering assigned eight spatial domains (layer I with leptomeninges, L1/pia; L2/3; L4; L5; L6; interneuron-rich; white matter; vascular/immune) that were concordant with the H&E anatomy and the laminar marker genes and occupied in both conditions (Fig. 6a to c; Supplementary Fig. 4). The control capture areas lie on gyral crowns and the NHD capture areas on the banks and fundi of sulci, so we did not compare laminar thickness or organization between conditions; in addition, the NHD white matter is sub-sulcal (U-fibre) white matter (Methods). Within each condition, the domains expressed their laminar markers, with some differences between conditions (Fig. 6b): in control, *RORB* increased from L2/3 to L5, *PCP4* peaked at L5 and *MBP* increased into white matter; in NHD, *CUX2* kept its superficial-to-deep gradient and *RORB* still increased toward L5, whereas the *PCP4* peak was lower and *MBP* was low in the white-matter domain, consistent with the under-representation of deep-layer neurons and the demyelination seen in the nuclei.

**Fig. 6.**
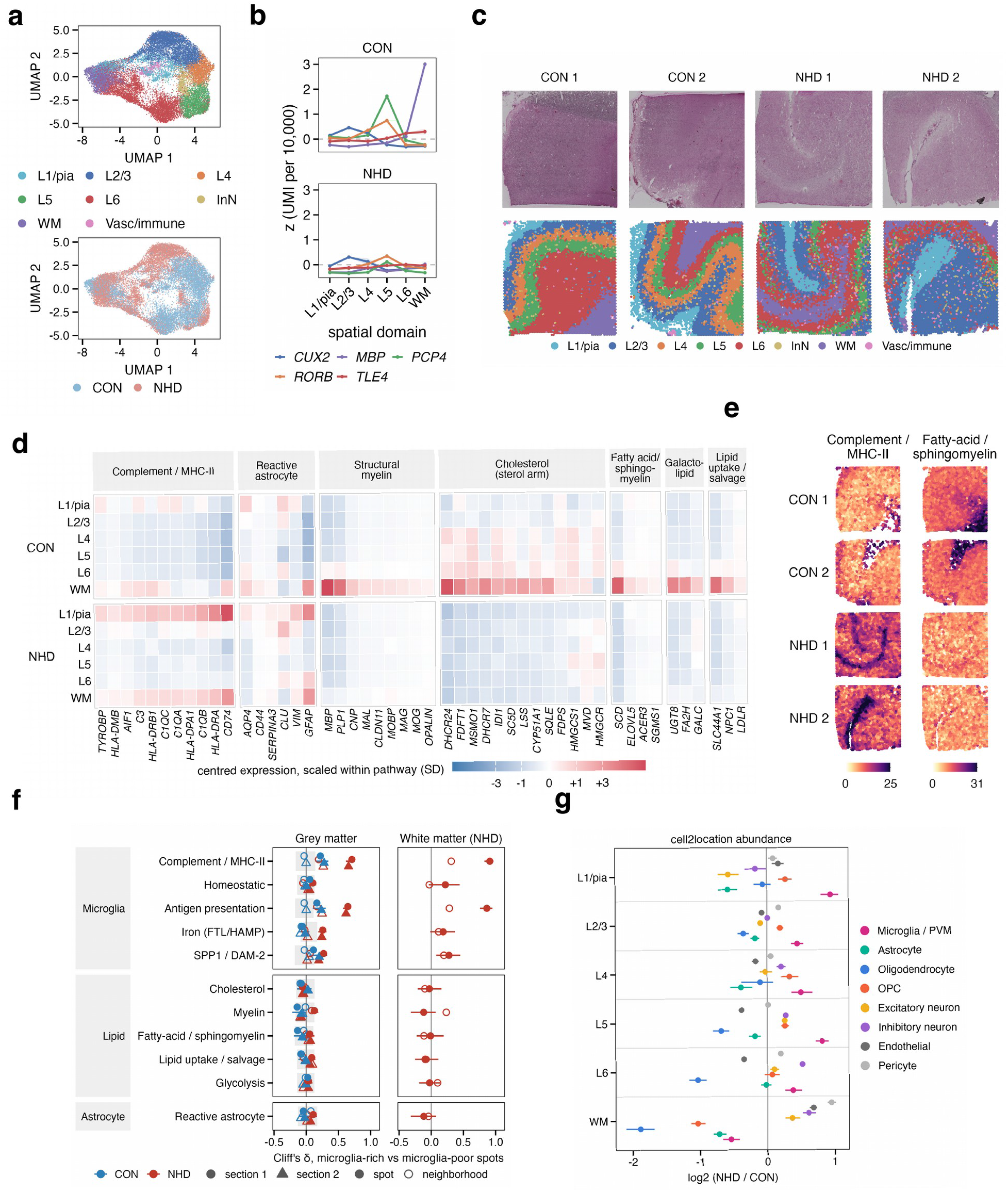
Spatial transcriptomics localizes the glial cascade in NHD prefrontal cortex. Two Visium sections per donor, 18,765 spots; values are per assigned domain, and no statistical test is applied (Methods). **a** UMAP embedding of all spots colored by spatial domain (top) and condition (bottom); eight domains from expression clusters plus a BANKSY-guided L1/pia domain (layer I with leptomeninges). **b** Laminar marker genes per domain within each condition (mean z of UMI per 10,000). **c** H&E image of each capture area above its domain map. **d** Heat map of program genes by domain and condition (mean UMI per 10,000, centered per gene and scaled within pathway). **e** The same programs on tissue for each section (UMI per 10,000, spot plus six neighbors). **f** Cliff’s delta between microglia-rich and microglia-poor spots, ranked within section and domain by their cell2location microglial fraction, for each program; circle, section 1; triangle, section 2; filled, own spot; open, six neighbors; bars, block-bootstrap 95% interval; grey band, toroidal null. Complement/MHC-II and antigen presentation are the positive controls for the ranking (Methods); white matter is NHD-only. **g** log2 ratio (NHD/CON) of the mean cell2location proportion of each cell type per domain; bars, block-bootstrap within-donor sampling intervals, not confidence intervals on a condition difference (all domains in Supplementary Data 11, sheet Composition per domain). Panel f uses depth-matched scores; b, d, and e native depth (Methods). Source data: Supplementary Data 11.

We first mapped the glial programs onto the domains as mean expression per spot (Fig. 6d, e). We found that they recovered the single-nucleus cascade *in situ*. Complement and MHC-II genes were elevated in every NHD domain, white matter included (*HLA-DRA* +1.32 to +1.90 log2, *CD74* +0.67 to +1.42, *C1QB* +0.85 to +1.40 across the five bands from L2/3 to white matter), and within NHD the program was concentrated in the two non-laminar compartments, layer I/pia and white matter, rather than in the layers (Supplementary Data 11, sheet Programs per domain). *GFAP* and *SERPINA3* were elevated in NHD grey matter and most in the upper layers, but not in white matter, where control expression was already high. The lipid supply chain ran the other way: the control white-matter domain expressed the myelin, cholesterol and sphingomyelin genes at 5- to 20-fold their grey-matter levels, each was lost in NHD white matter (*MBP* −2.81 log2, *PLP1* −2.68, *DHCR24* −2.74, *SCD* −3.67), and the fatty-acid/sphingomyelin program that marks the control white matter on tissue (21.1 per 10,000 against 2.7 to 4.9 in the control layers) was absent from the NHD white matter (2.3); within the layers the same arm decreased most in L5 and L6.

The cascade model predicts that the microglial state is local and its downstream programs are not. To test whether the programs are graded by local microglial content, we estimated the cell-type composition of every spot by cell2location deconvolution against the single-nucleus atlas, ranked the spots within each section and domain by their microglial fraction, and compared program expression between microglia-rich and microglia-poor spots as a Cliff’s delta, for each spot on its own and with its six neighbors (Fig. 6f; Methods). The complement/MHC-II and antigen-presentation programs, which serve as the positive control of the ranking because microglia express them, were concentrated in microglia-rich NHD spots (Cliff’s delta 0.68 [0.65, 0.71] and 0.63, against 0.25 and 0.20 in control spots ranked the same way), the iron and *SPP1* programs followed (0.25 and 0.23), and astrocyte reactivity and the cholesterol, myelin and lipid programs did not (|delta| ≤ 0.09, against a housekeeping module at 0.06); ranked by the six neighbors instead, complement/MHC-II and antigen presentation retained a gradient (0.23 and 0.21) and every other program decreased to |delta| ≤ 0.04, so at the 55-µm scale of a spot the microglial state is local and its downstream programs are not.

Composition on tissue was read from the same deconvolution, summed per domain (Fig. 6g). It placed microglia up in every NHD cortical layer (log2 NHD/CON +0.38 in L6 to +0.81 in L5) but not in white matter (−0.54), and oligodendrocytes down with cortical depth (−0.35 in L2/3, −0.69 in L5, −1.04 in L6) and most in white matter (−1.89), where OPC (−1.04) and astrocytes (−0.72) decreased as well. The spatial data thus recover *in situ* the directions the single-nucleus data give.

Across the cell-type programs, the same result held in two frames, effect size within the case and direction between prefrontal cortex and hippocampus, and, for the glial programs, in a third that does not depend on the shared donor, supporting a model in which DAP12 deficiency leaves microglia in a CIITA-driven, antigen-presenting state that does not engage the TREM2-dependent arm and is accompanied by a glial-to-neuronal degenerative cascade.

## Discussion

In this study we report the first single-nucleus and spatial map of the NHD brain across two regions, and we identified the state that microglia adopt when DAP12 is deficient. We found that DAP12 deficiency does not silence microglia: it leaves them in a CIITA-driven, MHC class II-high antigen-presenting state that does not engage the TREM2-dependent arm of the disease-associated trajectory. From that state the microglia remodel their secreted output onto astrocytes, oligodendrocytes and neurons, each of which changes in a direction that holds between prefrontal cortex and hippocampus; for the glia, those directions also agree with the occipital cortex of the Zhou et al. cohort^22^ independently of the donor the two studies share. Zhou et al. found that NHD microglia are dysregulated. We extend that work in four ways: we determined what the state consists of and which programs were depleted or dysregulated; we traced its secreted output to the main neighboring cell types; we investigated the same programs in intact tissue; and we found that the response to the NHD mutation also occurred in the hippocampus, a region that lies outside the frontal syndrome defining the disease.

We first defined the features of the NHD microglial state, which had not been resolved beyond a dysregulation signature. At the cell surface, MHC class II and the Fc receptors predominate, together with complement and scavenger receptors. The metabolism of these cells is glycolytic and iron-handling, and the phagocytic and lysosomal program increased in both regions (Fig. 2d) while the phagocytic receptors of the TREM2-dependent arm (*CLEC7A*, *ITGAX*) did not, so the receptor route into that program is the Fc receptors rather than TREM2. This is a phagocytic program rather than demonstrated phagocytosis: whether it achieves clearance is untested here, and *Tyrobp*- and *Trem2*-deficient mouse microglia fail to clear myelin debris^17^. The elevated program may therefore be a compensatory response that the cell cannot complete. The secretome, finally, comprises complement, ferritin, *GAS6* and the TNF-family cytokine *TNFSF13B*, with *SPP1* in the hippocampus. Two regulators may be responsible for this signature: CIITA, active in both regions, and STAT3, whose activity estimates were positive in both regions and reached FDR < 0.05 in the prefrontal cortex but not in the hippocampus.

Of these two regulators, CIITA raises the obvious question of what switches it on, since it is the transactivator through which interferon-γ induces MHC class II^32^. Interferon-γ itself, however, is not upregulated here, and the MHC class II genes and *CIITA* themselves carry the one enriched interferon-γ term. No type I interferon set was enriched in either region (Supplementary Data 5), and STAT1, the transducer that couples interferon-γ to CIITA, did not pass the regulon gate. What activates CIITA in the NHD brain therefore remains open.

The second regulator, STAT3, is the factor Zhou et al. placed at the center of the NHD microglial signature in occipital cortex, and it is also the factor that drives the oligodendrocyte stress response in CSF1R-related disorder^33^, so two microgliopathies and three datasets converge on it. The STAT3-related genes they reported, *IL4R*, *PTPN1*, *ITGAV*, *HAMP* and *CD163*, were upregulated in both of our regions (Supplementary Table S1), so their tissue-repair profile and our antigen-presenting profile are one state read through different gene sets. Zhou et al. also proposed that FcRγ-mediated signaling might compensate for loss of DAP12 in mice but not in humans. We find it engaged in the human brain as well: the Fc receptors were upregulated in both regions and dominate the surface of the NHD state. Whether that engagement restores DAP12-dependent function cannot be judged from transcripts, but the receptor repertoire argues against confining the route to the mouse. The state also resolves an apparent paradox: in a disease defined by loss of function, the microglia appear activated rather than at rest. It differs, however, from TREM2-null microglia derived from patient stem cells, which lose antigen presentation *in vitro*^24^. Three features could account for that difference: the *in vivo* stimulus environment, the intermediate amyloid pathology of this brain, and residual leaky initiation from the start-loss allele.

What the state lacks is equally informative. We found that the TREM2-dependent arm was not engaged in either region, and its absence has implications beyond the model in which it was defined. The two-step model was derived in mouse amyloidosis, where the second step brings *Lpl*, *Itgax*, *Clec7a* and *Lgals3* and requires TREM2^19,20^; in the NHD brain the entry genes changed in both regions while no endpoint gene passed the discovery gate in either. In prefrontal cortex, *CLEC7A* and *ITGAX* were detected with effect sizes near zero (Cliff’s delta +0.09 and −0.02), and *LPL*, *CST7*, *LGALS3* and *GPNMB* fell below the detection floor. In hippocampus, *CLEC7A* and *ITGAX* trended upward without reaching significance (Fig. 2e; Supplementary Table S1). This is the first evidence that the two steps are separable in a human brain: a genetic lesion of the adaptor left the first step intact in two regions and the second step unengaged at the discovery level, with only a sub-threshold trend in hippocampus, as the mouse model predicts for loss of TREM2-DAP12 signaling. The two-step architecture of the trajectory, therefore, holds in humans, and the entry step does not require DAP12.

The absent second step could also explain the Alzheimer-type change in NHD brains. The NHD donor carried Thal phase 2 amyloid and Braak III tau, and in Alzheimer’s disease with intact TREM2 that pathology recruits the lipid-handling program^34^; here it did not, and the 29-year course makes this donor the strongest available test of that expectation: a DAP12-deficient brain accumulated amyloid and tau across its sixth decade without recruiting the lipid program. The same logic makes Alzheimer-type lesions in NHD brains (this donor’s intermediate change at 59, the modest change reported across NHD brains^35^, and the substantial, atypically distributed tau pathology of a 51-year-old TREM2-null case^36^) a prediction of the model as well as an alternative explanation for the reactive-glial results: TREM2-dependent microglia compact plaques, limit their diffusion and impede tau seeding^37,38^, and microglia that have not engaged that arm cannot mount this response. The mouse results on complete loss, however, differ (haplodeficiency of *Trem2* or *Dap12* impaired the plaque barrier and worsened axonal dystrophy^37^, whereas complete *Tyrobp* deficiency was reported protective in one early amyloid model^39^), so the prediction is stated for the human disease. Whether the microglia reached this point along the trajectory and progressed no further, or never entered the TREM2-dependent arm, is a temporal question that a single end-stage time point cannot resolve; what the data show is a state, the same in both regions.

Why DAP12 deficiency produces this state, and why the disease is so much more severe than TREM2 hypofunction, may both follow from DAP12’s position as a hub. DAP12 couples TREM2, TREM1, SIRPβ1, CLEC5A and the activating CD300 receptors to SYK^9^, is required for CSF1R-driven proliferation^10^, and with FcRγ costimulates osteoclasts^11^; its loss abolishes signaling through all of these at once, which is why NHD affects bone as well as brain and why *TREM2* variants that raise Alzheimer’s risk do not cause NHD. Our data show the two sides of that hub in a single cell type. The DAP12-dependent route that we can assess, the TREM2 arm of the trajectory, is not engaged; and the other DAP12-coupled receptors are not expressed in human microglia at a level that can be assessed. The routes that are upregulated, by contrast, are those that do not require DAP12: *FCGR3A* and *FCGR1A* signal through FcRγ, and *FCGR2A* through its own ITAM; the adaptor transcripts themselves change little, and the CIITA/MHC class II arm is transcriptional rather than ITAM-dependent. We therefore interpret the NHD state as ITAM signaling redirected through the remaining receptors and adaptor proteins; this is a hypothesis at the transcript level, since SYK activity is regulated by phosphorylation, and TREM2 itself acts through SYK-dependent and SYK-independent routes^40^.

That hub is compromised in NHD even though the *TYROBP* transcript is present, and we therefore infer that the lesion in this donor acts at translation rather than at transcription. A start-loss substitution keeps the reading frame and introduces no premature termination codon, so the message escapes nonsense-mediated decay. In our data, it was not only retained but detected more often in NHD than in control microglia. This increase tracks the activation program, since *TYROBP* belongs to the disease-associated set. What this message cannot produce is DAP12 initiated at its own start codon. The mutant codon ACG is a near-cognate initiator that delivers methionine from the initiator tRNA, so any DAP12 made from it would be full-length and wild-type in sequence, at a fraction of the normal rate. Initiation at the next in-frame AUG (Met48) would not rescue a working protein: that codon lies in the transmembrane segment, and a protein begun there would lack the signal peptide and the extracellular cysteines through which DAP12 dimerizes. Reduced full-length protein is therefore the only product this allele can yield, and it has been detected: Sasaki et al.^7^ found abundant DAP12 in the microglia of an NHD brain with a homozygous single-base *TYROBP* substitution, confirmed by western blotting and quantitative PCR, against little or none in the other cases they examined. This is why the donor is DAP12-deficient rather than null, and why the transcript cannot serve as a readout of the lesion.

The Zhou et al. cohort^22^ allowed us to test whether this holds for the other NHD allele. Their three donors carry two kinds of allele: the start-loss allele of this study in two of them, ours among them, and in the third, c.141delG, an exon 3 frameshift for which loss of DAP12 protein has been reported^7^. They showed that *TYROBP was* confined to the microglial cluster, but did not report its levels by donor. In our re-analysis of their data, *TYROBP* was detected in the microglia of all three donors, in 21% and 16% of microglia for the two start-loss donors and in 43% for the frameshift donor, against 22% to 44% across their eleven controls, and not as a depth artifact, since the lowest NHD donor has deeper microglial nuclei than any control and the second lies within the control range (Supplementary Data 9, sheet TYROBP per donor). Neither NHD allele, then, removes the message. The frameshift transcript evidently escapes nonsense-mediated decay as well, so the two alleles differ in what can be translated rather than in whether the message survives: the start-loss allele can yield full-length DAP12 at a reduced rate, whereas the frameshift can yield only a truncated protein lacking the transmembrane and ITAM segments, consistent with the absent protein reported for it.

Whether the allele matters beyond the message is a separate question. Zhou et al. noted variability between their donors and raised the distinct mutations as one explanation. In our re-analysis the two start-loss donors resembled each other more than either resembled the frameshift donor in astrocytes, oligodendrocytes and neurons (excitatory neurons rho 0.45 against −0.02 and 0.17), though not in microglia (0.41 against 0.33 and 0.45; Supplementary Data 9, sheet Within Zhou donors); genotype also coincides with sex and postmortem interval in that cohort, so this is an observation and not an attribution. What the two studies do have in common is the microglial state itself: microglia adopt it in both regions of our donor and in their cohort with and without the shared donor, and this argues that the state follows from loss of DAP12 function rather than from the fate of its transcript. Whether residual full-length DAP12 from a start-loss allele modifies the disease is a question for protein-level work across genotypes, and NHD genotypes should be reported with their transcript and predicted protein fate.

With the state and its lesion defined, we next consider its effects on neighboring cell types. Because *TYROBP* was detected only in microglia among the cell types we profiled, the changes in astrocytes, oligodendrocytes and neurons are most likely non-cell-autonomous: the microglial secretome changed, and each neighboring cell type responded to it differently. We consider the three in turn. Zhou et al. modeled the microglia as receivers, with astrocyte-derived FGF2 predicted as the top ligand driving their signature; we model them as senders, and the two are complementary predictions from co-expression. Onto astrocytes the output was complement together with SPP1 and GAS6, secreted signals of the class through which activated microglia induce reactive astrocytes^16^, and the astrocytes became pan-reactive along a multidimensional axis^27,28^ rather than a neurotoxic pole. The wound-healing genes that Zhou et al. reported in NHD astrocytes (*F3*, *TNC*, *CD44*, *NRP1*) were upregulated here as well (Supplementary Table S1), with one program changing in the opposite direction: the metallothionein module — including *MT3*, the growth-inhibitory factor that buffers zinc and copper and restrains neurite outgrowth^29^ — was downregulated.

Beyond the astrocytes, the oligodendrocyte lineage is where the output reaches the tissue that defines the disease. Onto this lineage the trophic *TENM4*-*ADGRL3* pair was lost, *PDGFB*-*PDGFRA* gained in hippocampus and *PSAP*-*GPR37* was kept. The lineage itself changed in two ways that our analyses separate. Its composition changed: along the maturation continuum, mature oligodendrocytes were depleted and the remaining cells occupied two NHD-enriched states, and on tissue, deconvolved oligodendrocyte abundance decreased with cortical depth and most in white matter, where the myelin, cholesterol and sphingomyelin transcripts that mark control white matter were lost. And the surviving cells changed their program: they raised myelin transcripts while lowering cholesterol synthesis, a deficit that persisted along the continuum at matched depth. Zhou et al. reported reduced *MBP* and *ANLN* per nucleus in NHD oligodendrocytes of the occipital cortex; we reproduce the loss of *ANLN* and the upregulation of *KCNH8*, *CLIC4,* and *ADGRB3*, whereas *MBP* did not change per nucleus in our donor and was low only in the tissue, in the white matter, where oligodendrocytes are lost. At the same time, other myelin transcripts were upregulated (Supplementary Table S1). Whether this difference reflects region, disease stage, or analysis can be tested in their data. The same sterol-synthesis deficit appeared in prefrontal excitatory neurons, with the same trend in the hippocampus, so the loss of lipid synthesis is a property of the NHD brain rather than of a single lineage, with the oligodendrocyte lineage bearing its structural cost.

The program change requires a separate interpretation, because it separates two processes that a healthy oligodendrocyte runs as one. Myelin is about seventy percent lipid by dry weight, a quarter of it cholesterol, and oligodendrocytes synthesize most of that themselves: cholesterol synthesis is rate-limiting for myelin membrane growth^30^, and astrocyte-derived lipids only partly substitute when it fails^41^. A surviving NHD oligodendrocyte that raises *PLP1*, *MBP,* and *OPALIN,* while *SREBF2* and its targets decrease, has uncoupled the two. We interpret this as a compensatory response that cannot be completed: the surviving cells transcribe the myelin assembly program, carried by the *OPALIN*-high state, without producing the membrane lipid it requires. The microglial lesion may be what limits that lipid supply. Remyelination in the adult brain depends on cholesterol that microglia recover from myelin debris and pass to oligodendrocytes^42^, and dietary cholesterol accelerates repair when that supply is limited^43^. This handling requires TREM2, without which microglia fail to clear debris and to mount the cholesterol-transport program^17,44^; impaired cholesterol clearance by phagocytes is itself sufficient to limit remyelination^45^. DAP12-deficient microglia lack the TREM2 arm, so the lesion that leaves the microglia without the lipid-handling program would also leave the oligodendrocytes without the lipid that program supplies. The alternative interpretation, however, is feedback rather than starvation: *SREBF2*, *INSIG1,* and *LDLR* decrease together, as they do in a sterol-replete cell, and myelin debris in demyelinating white matter could be the source of that sterol. Transcripts cannot separate the two; a measurement of cholesterol in the tissue would. The compensatory interpretation is the one that makes a prediction: if the lipid supply were restored, the surviving oligodendrocytes could complete the myelin they have begun to transcribe.

The neuronal changes are the ones most closely associated with dementia. Onto neurons, the output was split by valence: a trophic GAS6 arm whose TAM receptors were downregulated, and a phagocytic SPP1 arm^18^ whose integrin receptors were downregulated, with the inhibitory purinergic pair lost. Excitatory neurons lost presynaptic and activity-dependent programs in both regions, which, in two further regions, confirms the reduced synaptic-transmission genes reported by Zhou et al. in the occipital cortex, and they lost oxidative phosphorylation in the prefrontal cortex. These are the energetically expensive functions that fail early in neurodegenerative diseases^46^, and both losses exceed what the shallower depth of NHD nuclei could account for.

Although it is challenging to determine which glial change contributes most to the neuronal deficit, our data allow us to rank the candidates. The microglial input to neurons changed, and its receptors were downregulated on deep-layer excitatory neurons; the astrocytic synapse-organizing input was preserved in the prefrontal cortex and decreased only in the hippocampus. The oligodendrocyte lineage offers a route that our ligand-receptor analysis does not capture: oligodendrocytes supply axons with lactate, and disrupting that support causes axonal degeneration^47,48^, so the loss of mature oligodendrocytes, which is greater in the prefrontal cortex, could contribute to the loss of oxidative phosphorylation in prefrontal excitatory neurons. We therefore regard microglial signaling and the loss of oligodendroglial support as the more likely contributors to the neuronal deficit, and astrocytic input as the less likely one in the prefrontal cortex, a ranking that would require experiments manipulating each cell type to test. The deep-layer excitatory subclasses were also under-represented among NHD nuclei; we keep this descriptive because nuclear composition is capture-biased, but it agrees with lower deep-layer marker expression in tissue and with the disease’s frontal predominance. A single microglial state thus projects three secreted profiles with three distinct consequences, thereby enabling a primary microgliopathy to reach the rest of the brain.

Our spatial data located these changes in the tissue, and every compositional claim of this study rests on them. Within the NHD grey matter, the microglial state concentrates in microglia-rich spots. However, astrocyte reactivity and the lipid programs are not graded by local microglial content at the 55-µm scale of a spot. Whatever carries the cascade from microglia to their neighbors, therefore, acts across the tissue rather than within individual spots, consistent with secreted signals rather than contact. We found that the deconvolved microglial fraction is higher in every cortical layer of NHD cortex, consistent with the preserved Iba1-positive microglia reported in NHD cortex by immunohistochemistry^23^, whereas in the sub-sulcal white matter it is lower alongside the depletion of oligodendrocytes and their precursors; that estimate is compositional, because the reference-inflated vascular fractions increase in the same domain, and it cannot test whether DAP12 deficiency depletes white-matter microglia as loss of CSF1R signaling does^10,33^. Glial composition differs most from control in the white matter. Within NHD tissue, the microglial program concentrates in the two non-laminar compartments, the layer I/pia of the sulcal walls and the sub-sulcal white matter, rather than in the cortical layers beneath layer I: the tissue that loses the most cells is also where the microglial state is most expressed.

Across all three receivers, the two regions responded differently to microglia in the same state. That difference is more plausibly imposed by the local environment than inherited by the microglia: adult homeostatic microglia are largely homogeneous across regions in mouse^49^, with strong regional states arising as niche subsets^50^, so a response that is region-restricted in the genes detected but shared in direction indicates one lesion expressed through different surroundings. In the hippocampus, the SPP1 input to astrocytes is larger, and astrocytes add a heat-shock arm to a broader reactive shift. In contrast, in the prefrontal cortex, the loss of mature oligodendrocytes is greater, with the loss of white-matter lipid programs observed *in situ*. The clinically central region is therefore not the one with the broader glial response but the one whose tissue is most dependent on the affected functions. The donor’s history offers a candidate for the hippocampal input: twelve years of epilepsy selectively injures the hippocampus, and we speculate that a seizure-injured niche drove the larger hippocampal microglial response to the same lesion; that reading is untestable in one donor and would be resolved by a second NHD hippocampus without seizures.

The white matter is where our study found the greatest effect. Myelin-clearing microglial states of white matter share their lipid-handling genes with the TREM2-dependent arm^51^, and TREM2-deficient mouse microglia fail to clear myelin debris and leave demyelination unrepaired^17^, so DAP12-deficient microglia would be expected to mount the inflammatory arm of the white-matter response without completing its clearance arm. Together, these give a reading of why a pan-microglial lesion yields a white-matter-predominant disease, stated as a prediction from a mouse-defined state.

Our approach has the strengths of a deeply characterized rare case and the limits of a single case. We had a genetically confirmed, neuropathologically staged donor with a sex- and ancestry-matched control from the same bank; two regions, dissected and processed independently, which allowed us to test each program for consistency within the donor; two modalities that read the same programs in nuclei and in tissue; and the Zhou et al. dataset, with two further NHD donors and independent controls, against which we could contrast every program with and without the shared donor. We have one NHD-control pair, so our inference rests on effect size, on coherence between regions, and on agreement with Zhou et al. rather than on single-donor significance^25^. The cross-cohort comparison separates the shared from the variable: the microglial and astrocytic programs agree with Zhou et al. independently of the shared donor, whereas the neuronal programs agree through that donor and vary between the other individuals and between occipital and frontal regions, as neuronal vulnerability does across neurodegenerative diseases; the neuronal cross-cohort agreement is therefore interpreted as consistency within our donor across regions and laboratories, and the glial agreement as the conserved core of the disease.

The limits of our design follow from the rarity of the material. In our donors, disease status is confounded with donor identity, postmortem interval, RNA integrity, Alzheimer-type change, and seizure history. The Alzheimer-type change is within the range of unselected sixth-decade brains^52^, has been described in NHD before, cannot set the direction of a cascade whose lesion is germline, and did not recruit the TREM2 arm; the seizure history is a competing explanation for hippocampal neuronal loss and, in both regions, for the loss of the immediate-early-gene program, while presynaptic and oxidative-phosphorylation programs are less exposed to it. RNA quality co-varies with the neuronal scores, but with seven samples in which integrity tracks condition, any condition effect does so as well, and the loss does not scale with the integrity gap between regions. Sequencing depth was modeled per comparison, which is why the hippocampal oxidative-phosphorylation and postsynaptic magnitudes are withheld, and hippocampal microglial magnitudes are upper bounds; the NHD hippocampus is one capture with the lowest RNA integrity, and the control hippocampal capture is interneuron-rich; and compositional claims rest on the spatial data, in which the control white matter is too shallow to enter the depth-matched comparisons. We infer cell-cell communication from co-expression; we did not partition astrocytes or microglia into substates at these numbers of nuclei; we use the mouse-derived DAM framework as a coordinate rather than an identity; and our conclusions are transcriptomic and spatial. Finally, we observe the endpoint of a lesion present from conception: the cells we profiled are those that survived nearly six decades of it, and the order in which the cell types changed is inferred rather than observed.

In conclusion, loss of DAP12 function leaves microglia in a CIITA-driven, antigen-presenting state that does not engage the TREM2-dependent arm. Through a remodeled secretome, that state is accompanied by astrocyte reactivity, loss of mature oligodendrocytes, and, in excitatory neurons, loss of presynaptic and activity-dependent programs and of prefrontal oxidative phosphorylation, most severely in the white matter. On this view, NHD is not a diffuse leukoencephalopathy but a stereotyped, non-cell-autonomous response with a single point of origin, whose glial arm is shared across the three NHD donors profiled to date. It suggests that, in the human brain, microglia do not merely respond to their environment but shape it. Because the same TREM2-DAP12 axis is graded in the common dementias (hypofunctional *TREM2* variants and a single deleted copy of *TYROBP* raise Alzheimer’s disease risk^13,14^, and partial loss of CSF1R produces a leukoencephalopathy of its own^33^), the NHD brain may show in complete form the microglial state that partial loss of this axis produces elsewhere. We propose that this state also points to where intervention could be effective in the common dementias: at steps downstream of TREM2, which TREM2-directed agents cannot reach without DAP12, or by restoring the trophic and cholesterol-synthesis support that fails in the cells surrounding microglia.

## Methods

### Human material

Postmortem tissue from two Japanese women was obtained through the Mihara Memorial Hospital Brain Bank (Isesaki, Japan) with consent for research use: one NHD donor (donor 861) and one control donor with no neurological diagnosis. Their clinical, genetic and neuropathological characteristics, taken from the Mihara records and neuropathology report (M.T.), are given in Table 1 and summarized in the Case presentation; occipital cortex from the NHD donor was profiled by Zhou et al.^22^. The NHD donor is homozygous for *TYROBP* c.2T>C (p.Met1Thr; NM_003332.4), a start-loss variant reported in Japanese NHD^6^ and classified pathogenic in ClinVar (VCV000005797), confirmed by Sanger sequencing. The variant introduces no premature termination codon, so *TYROBP* mRNA is expected to escape nonsense-mediated decay, which the retained transcript in NHD microglia confirms (Results); retained DAP12 protein has been reported for a homozygous single-base *TYROBP* substitution^7^, and DAP12 protein was not measured in this donor. We therefore describe the allele as start-loss, predicted loss of function, and the donor as DAP12-deficient. Postmortem interval, RNA integrity, intermediate Alzheimer-type change in the NHD donor^35,36,52^ and a twelve-year history of epilepsy differ between the donors and are reported per donor and sample (Table 1). The study was approved by the Keio University School of Medicine Ethics Committee (no. 20160273) and the RIKEN Yokohama Campus Ethics Committee (H30-26); written informed consent for the research use of the tissue was obtained from each donor’s next of kin through the brain bank.

### Study design

Two regions were sampled from each donor: prefrontal cortex (“Frontal” in figure panels and tables), dissected into grey-matter-enriched and white-matter-enriched fractions, and hippocampus, sampled whole. This yielded seven samples, each a nucleus preparation from one dissection, sequenced as its own library: four control libraries (frontal grey-matter-enriched, frontal white-matter-enriched, two hippocampal) and three NHD libraries (frontal grey-matter-enriched, frontal white-matter-enriched, one hippocampal). The prefrontal cortex from both donors was also profiled by 10x Visium, two sections each. The design has one donor per condition, which is the material this disease allows, and we built the inference to match it rather than to disguise it. Nuclei from one donor are not independent replicates of the disease, so a per-nucleus P value, however small, measures the separation of two cell populations from two individuals and cannot by itself show that a change belongs to NHD rather than to the donor^25^. We therefore use per-nucleus tests only to rank and nominate features, and we rest every claim on three lines of evidence that a single-donor P value does not provide. First, effect size: every reported quantity is an effect size (Cliff’s delta, NHD relative to control, for per-nucleus comparisons; log2 fold change; change in communication probability), measured on raw counts, with sequencing depth handled as described below, and gated at a pre-declared threshold. Second, consistency between regions: prefrontal cortex and hippocampus were dissected, captured, and sequenced as separate libraries, so a change that appears in both in the same direction has been observed twice in independent preparations of the same lesion, and agreement is judged against the rate expected by chance rather than against zero. Third, agreement with an external cohort: the Zhou et al. dataset^22^ provides two additional NHD donors, eleven independent controls, and a different region, laboratory, and pipeline, and we score directional agreement with and without the donor shared by the two studies. A change that clears the effect-size gate, holds between regions, and agrees with the external cohort has passed three tests that a donor artifact would have to pass together; a change that clears only the first is reported as such. The only inferential mark on any panel is a grey q*, which flags a comparison that reached adjusted P < 0.05 without reaching the effect-size gate (|Cliff’s delta| < 0.15), so that a reader can see where significance and effect size disagree. A discovery feature is a gene with Bonferroni-adjusted P < 0.05, |Cliff’s delta| ≥ 0.15, and detection ≥ 10% on raw counts after the artifact filter described below.

### Single-nucleus RNA sequencing

Nuclei were isolated from two 150-µm frozen sections per sample with the Chromium Nuclei Isolation Kit (10x Genomics) according to the manufacturer’s instructions, stained with 7-aminoactinomycin D (7-AAD), and 7-AAD-positive nuclei were purified on a MoFlo cell sorter (Beckman Coulter). Purified nuclei were processed with the Chromium Next GEM Single Cell 5′ Reagent Kit v2 (10x Genomics), and libraries were sequenced in a single batch on a NovaSeq X Plus (Illumina) with 150-bp paired-end reads (Supplementary Data 2, sheet QC per sample). Reads were aligned with Cell Ranger 7.1.0 against GRCh38-2020-A; ambient RNA and empty droplets were removed with CellBender 0.3.2^53^ (false-positive rate 0.01, 100 epochs, expected cells set to the Cell Ranger call and total droplets to 1.2 times that value); multiplets were removed with scDblFinder. We retained nuclei with 200 to 6,500 detected genes and mitochondrial content below 10%, leaving 55,354 of 64,162 Cell Ranger-called nuclei (13.7% removed; control 37,371, NHD 17,983; frontal 37,034, hippocampus 18,320; 3,624 to 12,309 per library; Supplementary Data 2, sheets Nucleus counts and QC per sample). A T-lymphocyte class (60 nuclei) was annotated separately, retained in the dataset, and excluded from the myeloid analyses; it is shown beside the myeloid nuclei only in the subclustering of Supplementary Fig. 3d. Within Micro-PVM (3,081), 95 oligodendrocyte doublets were removed at re-clustering. The CD163^+^/F13A1^+^ cluster (162) was held out, leaving 2,824 microglia for the microglia-only analyses (Supplementary Data 2, sheet Myeloid selection).

### Annotation

Nuclei were processed in Seurat v5^54^, normalized per library with SCTransform v2^55^, integrated over library with Harmony^56^ on 30 principal components, and clustered at resolution 1.0. We assigned broad labels via Azimuth transfer against the human motor cortex reference, collapsed to eight classes: Micro-PVM (microglia with perivascular macrophages, the reference’s class name), astrocytes, oligodendrocytes, oligodendrocyte precursor cells (OPC), excitatory neurons, inhibitory neurons, endothelial cells, and pericytes. Within Micro-PVM, a CD163^+^/F13A1^+^ population was resolved and reported by its markers, with no origin asserted; the pooled class was used for gene-level and cell-cell communication analyses, microglia alone for program scores, and the myeloid compartment for regulon activity (Supplementary Data 8, sheet Pooled vs microglia-only). Frontal excitatory neurons were annotated by Seurat label transfer (3,000 variable features, 30 principal components; 9,653 anchors) against the dorsolateral prefrontal cortex reference of Jorstad et al.^57^, using its excitatory nuclei with each subclass capped at 6,000 (31,591 nuclei); this reference resolves the nine excitatory subclasses of granular cortex, including L4 IT. Transfer confidence was equal between conditions (median prediction score 1.00 in both). Inhibitory neurons were annotated with Azimuth, giving 15 cortical subclasses (Supplementary Data 2, sheet Frontal subclasses). Subclass shares depend on dissection depth and on the capture bias of nuclear preparations, so they are reported as descriptive, matter-matched values (grey against grey, white against white). Hippocampal neurons were subclustered and each cluster scored against a 60-marker human-hippocampus panel^58,59^, with every call required to carry at least one marker private to its label at ≥ 10% detection. This resolved a pyramidal cluster carrying CA3 markers (*HS3ST4*, *GRIK4*, *NECAB1*), which likely also contains CA1 and CA2 nuclei, subiculum, and CGE (VIP, LAMP5) and MGE interneurons (Supplementary Data 2, sheet Hippocampal subtypes); one cluster of 190 nuclei with no private marker, the lowest complexity and 186 nuclei from one sample is reported as unresolved and excluded from subtype comparisons. Dentate granule and Cajal-Retzius neurons were not recovered, so hippocampal composition is not interpreted. Oligodendrocytes were subclustered as a maturation continuum with two further states, high-myelin and stress, each required to be NHD-enriched and to score on a curated module (so their NHD predominance is in part definitional); OPC were analyzed separately.

### Sequencing depth

Sequencing depth differs between libraries and between conditions within a cell type, and a shallower nucleus detects fewer genes and scores lower on any program, so depth could imitate a loss. We handled it in three ways, so that no result depends on libraries being equally deep: counts were library-size normalized for every effect size; log10(nCount_RNA) entered every per-nucleus MAST model as a covariate (Supplementary Data 3); and, for each program, we measured how much change depth alone can produce and withheld any change that did not exceed it. Because the NHD-to-control ratio of median depth varies by compartment — NHD nuclei are deeper in hippocampal microglia (3.4) and OPC (1.6) and shallower in frontal oligodendrocytes (0.54), frontal astrocytes (0.66) and hippocampal excitatory neurons (0.22; Supplementary Data 10) — depth was assessed comparison by comparison. Where depth differs between conditions in the same direction as a change, the change was tested against the largest effect depth alone could produce: control nuclei were split into deep and shallow strata at the observed NHD-to-control depth ratio, the program was scored in each stratum, and the resulting delta, which reflects depth and nothing else, is the bound; program changes within that bound (hippocampal oxidative phosphorylation, −0.47 against 0.53; postsynaptic, −0.17 and −0.28 against 0.44 and 0.47) are not reported as effect sizes. NHD hippocampal microglia carry 3.4 times the median RNA content of control hippocampal microglia, an increase confined to microglia and not explained by ambient RNA or doublets, so hippocampal microglial magnitudes are upper bounds. Down-sampling every nucleus to 750 unique molecular identifiers was used as a sensitivity test and as the estimator for the oligodendrocyte pseudotime curves (Fig. 4i).

### Differential expression

Differential expression was run in two tiers, because the two available units of analysis answer different questions: the nucleus, which gives power to detect a change but treats every nucleus as a replicate, and the library, which respects the sample structure but gives only two libraries per condition in prefrontal cortex. In the discovery tier, NHD-versus-control differential expression was computed per cell type and region with MAST^60^ on the SCT assay (genes detected in at least 10% of nuclei, minimum log fold change 0.1), with dissection block and log10(nCount_RNA) as covariates, and adjusted by Bonferroni over all 26,973 assay features. Effect size and detection were measured on the raw counts: a discovery feature has adjusted P < 0.05, |Cliff’s delta| ≥ 0.15^61^ and detection ≥ 10% in at least one condition (Supplementary Data 3). A fixed blocklist, applied identically to every cell type, removed neuronal transcripts scored in glia, unannotated lncRNA and antisense classes, histones, housekeeping, sex-chromosome and unnamed genes (611 of 5,827 features); Supplementary Data 3 retains the unfiltered set. In the confirmation tier, counts were summed to library-level pseudobulk and modeled in DESeq2^62^ with apeglm-shrunken effect sizes, and a feature was called confirmed when it met adjusted P < 0.01, |log2 fold change| > log2(2.5) and agreed in sign with MAST. Prefrontal cortex, with a grey-matter-enriched and a white-matter-enriched sample for each donor, supports this tier; for the hippocampus, with one NHD sample, the pseudobulk model provides an aggregate shrunken fold change only, and a hippocampal feature is called supported when its direction agrees with the prefrontal pseudobulk estimate (Supplementary Data 4).

### Enrichment, signature scores and regulon activity

Gene-set enrichment used fgsea^63^ on genes ranked by log2 fold change trimmed at the 1st and 99th percentiles, against GO biological process, Reactome and Hallmark and, for microglia, 19 published state signatures, with Benjamini-Hochberg correction within each comparison (Supplementary Data 5); the genome-wide microglial pathway view (Supplementary Fig. 3b) restricts the ranking to genes detected in at least 10% of nuclei in both conditions and free of ambient neuronal markers. Signature scores were computed per nucleus with AddModuleScore, or as the mean of normalized expression for the neuronal panels, and compared between conditions by two-sided Wilcoxon test with Benjamini-Hochberg correction and an effect-size gate of |Cliff’s delta| ≥ 0.15; gene sets, symbol mapping and any exclusion are listed in Supplementary Data 6, sheet Signatures (four of the ten genes of the mouse-derived complement/interferon-reactive set have no human ortholog and were not scored). Because glycolytic genes also increase with hypoxia in agonal tissue, the microglial glycolytic program was checked against the hypoxia-response gene set, which was depleted in NHD neurons in both regions (FDR < 0.001) and not enriched in the other glia (Supplementary Data 5), so the glycolytic gain is confined to microglia. Transcription-factor activity (Fig. 2h) was inferred with decoupleR^64^ on the signed CollecTRI regulon^65^ from per-gene Cliff’s delta across 2,985 of the 2,986 myeloid nuclei (one lacked a regulon score), with Benjamini-Hochberg correction over 118 factor-by-region tests and a pre-declared six-criterion gate (coverage, cross-region reproducibility, robustness to removal of housekeeping genes, agreement with the factor’s own transcript, mode-of-regulation purity, canonical anchors); SPI1 served as an identity control and PPARG as a declared null, and scores were stable to down-sampling at 750 unique molecular identifiers (Supplementary Data 8, sheet TF activity Fig2h).

### Cross-cohort comparison

We contrasted effect direction with the Zhou et al. cohort^22^: raw feature-barcode matrices from GEO GSE190015 (3 NHD, 11 controls, occipital cortex) reprocessed through our pipeline to 63,358 nuclei and re-annotated to matching classes (Supplementary Fig. 2). For each cell type and region we report the fraction of genes significant in both datasets (ours under the discovery gate, theirs at adjusted P < 0.10) that agree in direction, the empirical baseline agreement over all shared genes, and Spearman correlation over the shared set; significance is a one-sided binomial test against the baseline, and because genes are not independent the P values are exploratory (Supplementary Data 9, sheet Direction consistency). Their cohort comprises occipital cortex from three NHD donors — ours and a second homozygous for the same c.2T>C variant, and a third carrying a homozygous *TYROBP* deletion (c.141delG; given as c.141Gdel in that study) —with eleven independent controls (donor-arm correlations use Zhou et al.’s unshrunken pseudobulk effects, whereas the panel-level correlations of Figs 3c and 5c use apeglm-shrunken effects; both are in Supplementary Data 9, sheet Donor arms); among the two non-shared donors, genotype is confounded with donor identity and sex, so individual and regional contributions to the neuronal agreement cannot be separated. Because that cohort contains our donor, we also recomputed their side with the NHD group restricted by donor — the two donors not shared with this study, each other leave-one-out pair, and each donor alone — against the same controls, holding our effects fixed (Supplementary Fig. 2e; Supplementary Data 9, sheet Donor arms). Sex-linked genes do not contribute to the agreement: the discovery contrast is female versus female, and removing them left the statistics unchanged.

### Cell-cell communication and trajectory

We inferred ligand-receptor signaling with CellChat v2^66^ per condition and region (10%-trimmed mean, minimum 10 cells) in a glial run and a neuronal run; because labels are permuted within one donor per condition, changes in communication probability are interpreted as effect sizes only. In the astrocyte-to-neuron panel, pairs whose astrocytic ligand is on the ambient-neuronal artifact list (*NRXN1*, *NRG3*) were excluded, since their astrocytic signal cannot be separated from ambient neuronal RNA. Pseudotime was computed with Monocle 3^67^ on the oligodendrocyte object, with nuclei ordered from the highest early-marker score.

### Spatial transcriptomics

Ten-micrometre fresh-frozen sections for Visium were cut from the same prefrontal tissue blocks used for the single-nucleus libraries. Visium libraries (slide V11T10-025) were processed with Space Ranger 1.3.1 against GRCh38-2020-A. For section NHD 2, in which the fiducial alignment placed the barcode layout in mirror image relative to the scan, barcode positions were reflected about the array center (the transform leaving the fewest UMI-rich barcodes on background), and registration of every section was checked against the H&E (Fig. 6c). Tissue was called per barcode from the Space Ranger call supplemented by an H&E-mask tier that admits barcodes on stained tissue containing cortical neuropil (*SNAP25*, *RBFOX3* and *SLC17A7* ≥ 12 UMI per 10,000), so that the pial rim is included; 14 NHD 2 barcodes under an exogenous particulate contaminant were excluded. Spots were retained above a condition-blind floor of 50 unique molecular identifiers and 25 genes with no mitochondrial ceiling (percent mitochondrial reads regressed out), giving 18,765 spots (control 9,359, NHD 9,406). We segmented domains in Seurat with Harmony over section (30 principal components, resolution 0.5), sub-clustering the control upper-cortex cluster into a *CUX2*/*LAMP5*-high L2/3 band and a *RORB*-high L4 band. Because layer I shares expression features with white matter in the NHD sections, the L1/pia domain (layer I with leptomeninges) was assigned from neighborhood-aware BANKSY pial clusters^68^ (λ = 0.2, six neighbors) within the layer-I band annotated on the H&E of each section, blind to the transcriptome (1,039 spots; a histology-only alternative rule is carried per spot in Supplementary Data 11, sheet Spot annotation); the ten expression clusters and the BANKSY-guided L1/pia id were collapsed to eight domains by their highest-scoring laminar marker, with white matter and vascular/immune assigned by tissue position (Supplementary Fig. 4; Supplementary Data 11, sheet Spot annotation). The control capture areas lie on gyral crowns and the NHD capture areas in sulci, so comparisons are made per assigned domain and laminar thickness and order are not compared; domain occupancy ranges from 4% to 79% NHD spots and correlates with depth (Spearman rho −0.57), and no statistical test is applied to any spatial panel, because spots are sub-sampling units of one donor per condition. Program scores (AddModuleScore) were computed on a depth-matched matrix (spots ≥ 3,000 unique molecular identifiers, down-sampled to that depth; 14,804 spots; one of 976 control white-matter spots qualified, so white-matter comparisons are within NHD), whereas the laminar profiles, gene heatmap and program maps of Fig. 6b, d and e use all spots at native depth as gene or program UMI per 10,000, the maps pooled over each spot and its six neighbors. Cell-type abundance was estimated with cell2location^69^ against a reference regression fitted on an earlier eight-class single-nucleus dataset (61,256 nuclei, subsampled to 18,169) and summarized per domain as log2(NHD/CON) of the mean proportion with block-bootstrap 95% intervals (6 × 12 array blocks within a section, 2,000 draws); these are within-donor sampling intervals. In white matter the microglial ratio (−0.54) decreases as the reference-inflated vascular fractions increase and is read as compositional; in L4 the two NHD sections disagree on the oligodendrocyte ratio (−0.68 and +0.03). For the proximity analysis, spots of each section were ranked within each domain by their cell2location microglial fraction, or by the mean fraction of their six neighbors, and Cliff’s delta compared the depth-matched program scores of the top and bottom quartiles, with block-bootstrap intervals and a null from 200 toroidal shifts of the ranking map; the complement/MHC-II and antigen-presentation programs, whose genes are among the most microglia-specific in the deconvolution reference, serve as the positive control; grey-matter domains including L1/pia were pooled, and white matter was analyzed in NHD section 1, section 2 having fewer than 60 depth-matched white-matter spots once its layer I was assigned to L1/pia.

### Statistics and reproducibility

#### Design and replication

The study comprises one NHD donor and one control donor, the tissue available for a disease with about 200 reported cases. No randomization or blinding applies to a one-versus-one postmortem design. Disease status is therefore confounded with donor identity, postmortem interval, RNA integrity, Alzheimer-type change and seizure history, and the framework described under Study design is how we address that confounding: no claim rests on a single-donor P value, every claim is an effect size, and every claim is tested for consistency between two independently prepared regions and for agreement with an external cohort that includes two other NHD donors. Where one of these variables could affect a specific result, we tested it directly. The neuronal program losses did not scale with the RNA-integrity gap between regions (Supplementary Data 7, sheet RIN per library), and excluding the 189 unresolved hippocampal nuclei changed the reported deltas by at most 0.08 (Supplementary Data 7, sheet Unresolved sensitivity). Seizure-related injury remains a competing explanation for the absence of dentate granule neurons, for part of the hippocampal presynaptic loss and for the immediate-early-gene loss, so those results are read on direction and cross-region agreement rather than on magnitude.

#### Tests and effect sizes

Per-nucleus comparisons used two-sided tests, the MAST hurdle model for genes and the Wilcoxon rank-sum test for program scores, with the multiple-testing adjustment stated for each analysis: Bonferroni over all 26,973 assay features for the discovery table (Supplementary Data 3), and Benjamini-Hochberg within each panel or comparison elsewhere, with the family size given in each legend (14 tests in Fig. 2d, 22 in Fig. 3e,f (eleven signatures in two regions), 32 in Fig. 4d and 20 in Fig. 5d). Each comparison is reported with n nuclei, the exact adjusted P where it lies above the double-precision floor, and Cliff’s delta as the effect size. We chose Cliff’s delta because it is a rank statistic, bounded between −1 and 1, invariant to monotone transformations such as the log, and readable as the probability that a random NHD nucleus exceeds a random control nucleus minus the reverse. It is bounded by detection: when a gene is detected in a minority of nuclei in both conditions, the attainable |delta| is small (about 0.11 for *CLEC7A* in prefrontal microglia), so a null delta at sparse detection is reported against that ceiling rather than as evidence of no change. Pseudobulk comparisons used DESeq2 Wald tests with apeglm-shrunken effect sizes; cross-cohort agreement used one-sided binomial tests against each comparison’s empirical rate of chance agreement; and no statistical test was applied to the spatial data, in which spots are sub-sampling units of one donor per condition.

#### Held-out sets

Beyond the quality-control thresholds stated above, the sets held out of specific analyses are named where they arise: 95 oligodendrocyte doublets and 162 CD163^+^/F13A1^+^ nuclei from the microglia-only analyses, 60 T lymphocytes from the myeloid analyses, the 190-nucleus unresolved hippocampal cluster from subtype comparisons, the L5 ET and Sst Chodl subclasses and three subtype-by-region cells with fewer than 20 nuclei per condition from subtype tests, 611 artifact features from the discovery set, and 14 Visium barcodes under a contaminant. Detection gates for program scores (three set members, or two covering half the set, detected in 10% of nuclei) and the depth bounds are stated in the legends of the panels they apply to, and the gate outcome for every program-by-region cell is carried in Supplementary Data 7, sheet Program deltas.

#### Reproducibility

All analyses are scripted, with the random seeds listed under Software; the code and the per-panel source tables are released (Code availability), and every reported quantity can be traced to a row of the Supplementary Data.

### Software

R 4.5.1 with Seurat 5.0.1.9001, harmony 1.2.3, MAST 1.33.0, DESeq2 1.48.2, apeglm 1.30.0, fgsea 1.34.2, decoupleR 2.14.0, CellChat 2.2.0.9001, Monocle 3 1.4.27, Banksy 1.4.0, sctransform 0.4.2, msigdbr 26.1.0, Azimuth 0.5.0, glmGamPoi 1.20.0 and scDblFinder 1.22.0; Cell Ranger 7.1.0, Space Ranger 1.3.1, CellBender 0.3.2; cell2location 0.1.5 and scvi-tools 1.3.3 under Python 3.10, with image handling in Python 3.11. Random seeds: fgsea 1,000 simple permutations with sets of 10 to 500 genes (9,197 sets per comparison; seed 42 for the detection-gated run), Monocle 3 seed 42, sub-clustering seed 1,000, block bootstraps and toroidal shifts seed 42, and the split-UMAP down-sampling seed 42.

## Supporting information

Supplementary Data index

Supplementary Data 1: Data dictionary

Supplementary Data 2: Samples, nuclei and annotation

Supplementary Data 3: Per-nucleus differential expression (MAST)

Supplementary Data 4: Pseudobulk differential expression (DESeq2)

Supplementary Data 5: Gene-set enrichment

Supplementary Data 6: Gene sets and markers

Supplementary Data 7: Program effect sizes and sensitivity

Supplementary Data 8: Microglia

Supplementary Data 9: Cross-cohort contrast with Zhou et al.

Supplementary Data 10: Sequencing-depth diagnostics

Supplementary Data 11: Visium spatial transcriptomics

Supplementary Data 12: Cell-cell communication and composition

## Data availability

Processed count matrices, per-nucleus and per-spot metadata and Space Ranger outputs for all 55,354 nuclei and four Visium sections, together with the pipeline outputs, are available at Synapse under project syn77518970 (doi:10.7303/syn77518970, activated upon publication); the project is accessible to editors and referees on request during review and the processed data will be released upon publication. Raw sequencing reads, being human genotype-bearing data, are held under a Synapse controlled-access requirement and released to qualified researchers on approval by the Synapse Access and Compliance Team. The published NHD cohort of Zhou et al. 2023 was obtained as raw feature-barcode matrices from the Gene Expression Omnibus (GSE190015) and reprocessed through the pipeline described above. The dorsolateral prefrontal cortex reference of Jorstad et al.^57^ was obtained from the CELLxGENE collection d17249d2-0e6e-4500-abb8-e6c93fa1ac6f. For each Visium capture area the deposition comprises the Space Ranger raw and filtered count matrices, the spatial folder (H&E images, scale factors, barcode positions) and a spot-annotation table with the spatial domain and cell2location abundances used in Fig. 6; for section NHD 2 the deposited barcode positions and tissue call are those used in the analysis (Methods), so that the filtered matrix corresponds to the 4,598 analyzed spots while the raw matrix is the unmodified Space Ranger output. A data dictionary that defines every column of every sheet of Supplementary Data 2 to 12 is provided as Supplementary Data 1.

## Code availability

The analysis code (R and Python scripts, with package versions) is available at https://github.com/JulioLeonIncio/NHD-frontal-hippocampus-snRNAseq-Visium (tag v1.1-submission) and archived at Zenodo (https://doi.org/10.5281/zenodo.23112084).

## Acknowledgements

This publication is part of the Single Cell Medical Network of Japan. This research was supported by a research grant to the RIKEN Center for Integrative Medical Sciences (IMS) from the Ministry of Education, Culture, Sports, Science and Technology (MEXT) and a RIKEN Center Director’s Discretionary Budget Grant to J.L. J.L. is supported by the Alzheimer’s Association (award AARFD-24-1308438). M.T. is supported by the Japan Agency for Medical Research and Development (AMED; grants 21wm0425019 and 25wm0625126), the Intramural Research Grant for Neurological and Psychiatric Disorders of NCNP (6-8) and JSPS KAKENHI (22H04923).

## Author contributions

J.L. and S.M. conceived the project and designed the research. S.M., J.S., P.C., C.C.H., K.K., C.C. and H.O. secured funding and provided resources and institutional support. Y.A. coordinated the project. M.T. provided the human brain samples and performed the autopsies. I.K., S.Y. and J.L. processed the tissue and performed experiments. J.L. performed all bioinformatic analyses, prepared and assembled all figures and tables, and wrote the first manuscript draft. G.P. and J.M. performed data analysis. J.L. and S.M. led the project, interpreted the results and revised the manuscript with input from all co-authors. All authors read and approved the final manuscript.

