## Supplementary Information: Supplementary Figures 1 to 5, Supplementary Table S1 and Supplementary Data legends for "DAP12-deficient microglia adopt a TREM2-independent antigen-presenting state with a glial-to-neuronal cascade in Nasu-Hakola disease"

Contents: Supplementary Figs. 1 to 5 with legends; Supplementary Table S1; legends of Supplementary Data 1 to 12 (provided as separate workbooks).

### Supplementary figures

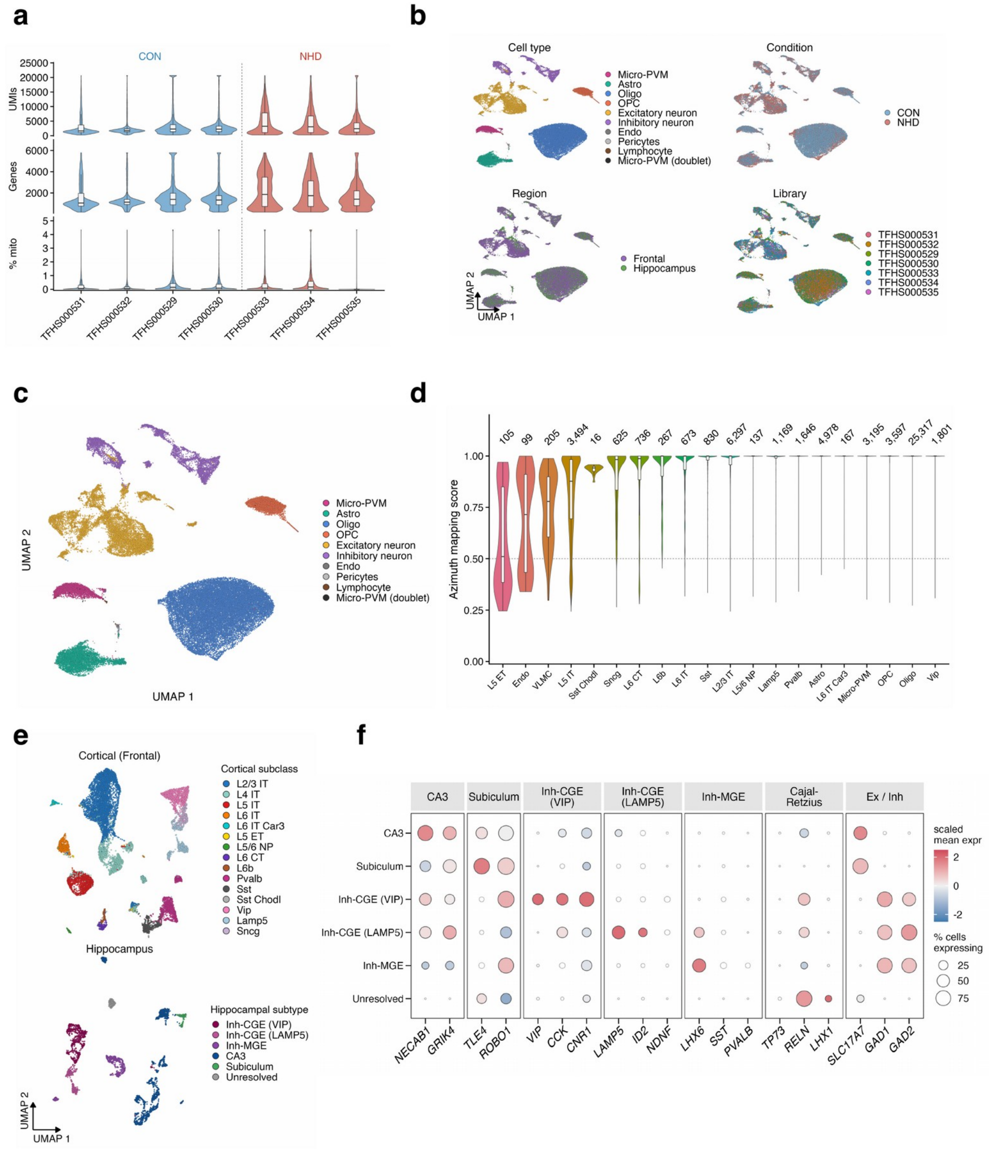


**Supplementary Fig. 1 Single-nucleus quality control, annotation, and hippocampal neuronal taxonomy. a** Per-library UMIs, detected genes, and mitochondrial fraction, shown to the 99th percentile. The NHD frontal libraries have more UMIs per nucleus (median 3,230 and 3,040, against 1,632 and 1,783 in control), a whole-library value that reflects their neuron-rich composition rather than greater depth within cell types. Depth enters every per-nucleus model as a covariate and is handled separately for each comparison (Methods, Sequencing depth). Mitochondrial content is at most 9.95%. RIN per library: NHD 6.3, 6.0, and 4.4; CON 7.4, 7.5, 7.1, and 6.7. **b** Embedding colored by cell type, condition, region, and library. Nuclei group by cell type and intermix across the other three variables. **c** Eight classes: Micro-PVM 3,081, astrocytes 4,978, oligodendrocytes 25,317, OPC 3,597, excitatory neurons 11,876, inhibitory neurons 6,087, endothelial cells 99, and pericytes 205, plus a small T-lymphocyte cluster (60) excluded from downstream analyses. **d** Azimuth mapping score per returned label (20 labels, all nuclei; median 1.000, with 98.5% of nuclei at or above 0.5). The counts above each violin are the Azimuth labels before hippocampal re-annotation; for Micro-PVM they include the lymphocyte cluster. **e** Cortical subclasses (frontal; 15, with excitatory labels transferred from a prefrontal reference and inhibitory labels from Azimuth) and five hippocampal subtypes defined *de novo*, beside one unresolved cluster (grey; 190 nuclei) that was excluded from subtype-level comparisons. The unresolved cluster is the lowest-complexity cluster in the hippocampus (median 1,312 genes, against 2,543 to 5,458 in the others). Dentate granule and Cajal-Retzius neurons were not recovered, and CA1/CA2 did not separate from the CA3-marker cluster, so interneurons make up 48.9% of the resolved hippocampal neurons. **f** Hippocampal subtype markers, including a Cajal-Retzius block, with the class markers *SLC17A7* (excitatory) and *GAD1* and *GAD2* (inhibitory). Dots for genes detected in fewer than 10% of a group's nuclei are hollow; *TP73* is hollow in every column. Dot size, percentage of nuclei; fill, scaled mean expression. The panels are descriptive, with no statistical comparison. Source data: Table 1; Supplementary Data 2 and 6.

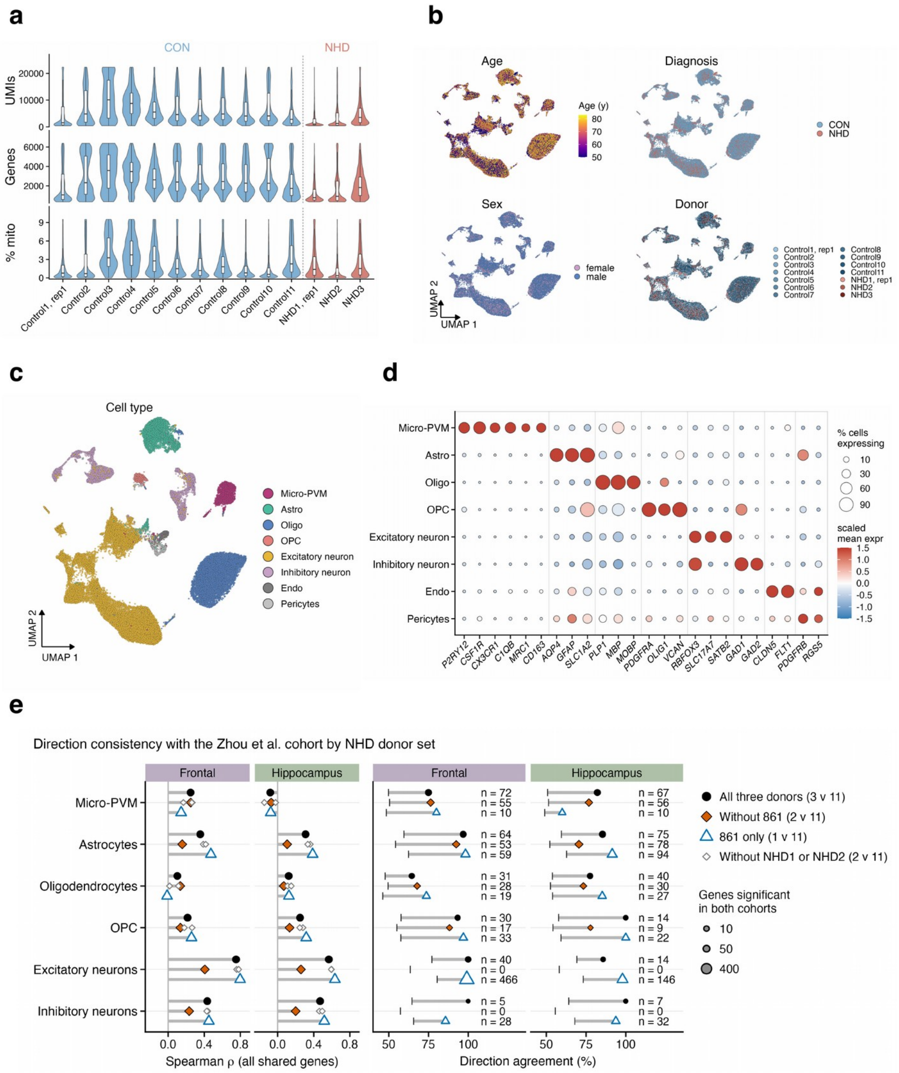


**Supplementary Fig. 2 Re-processing of the Zhou et al. cohort and cross-cohort agreement by NHD donor set.** Occipital cortex from 11 control and 3 NHD donors, 63,358 nuclei (CON 49,065, NHD 14,293); our NHD donor is one of the three. **a** Per-donor UMIs, detected genes, and mitochondrial fraction. Median depth is 4,070 UMIs in control and 1,523 in NHD, and batch and diagnosis are nearly confounded: two of the three NHD donors and the lowest-depth control share one batch. **b** Harmony embedding colored by age, diagnosis, sex, and donor (45,000 nuclei shown). No cluster is private to one donor or diagnosis. **c** Eight classes: Micro-PVM 2,884, astrocytes 6,053, oligodendrocytes 12,888, OPC 1,389, excitatory 30,693, inhibitory 7,738, endothelial 656, and pericytes 1,057. **d** Canonical lineage markers used for the annotation (fill capped at ±1.5). **e** Agreement with the Zhou et al. cohort by NHD donor set, for all six cell types and both regions. The NHD side of the Zhou et al. cohort is restricted by donor and compared against the same controls (11; 10 for Micro-PVM): all three donors (black circles), the two donors not shared with this study (vermilion diamonds), each other leave-one-donor-out pair (small grey diamonds, with the same power as the primary contrast), and donor 861 alone (blue open triangles; one NHD sample with dispersion estimated from the controls, descriptive only). Our per-nucleus effects are the same in every arm. Left, Spearman rho between our effects and the unshrunken pseudobulk effects of Zhou et al. over all shared genes. Right, the percentage of genes agreeing in direction among those significant in both cohorts, with n printed; the bar marks the genome-wide sign-agreement baseline, and "n = 0" marks arms in which no gene is significant in Zhou et al. Prefrontal microglial agreement is unchanged without donor 861 (rho 0.25 in both arms). Astrocytic direction agreement persists at a lower rho (92% of 53 genes, baseline 54%; rho 0.36 to 0.16). The excitatory-neuron correlation decreases (0.75 to 0.41), with no gene significant in both cohorts, whereas the other two leave-one-out sets stay at 0.76 to 0.78. The glial agreement is therefore shared across donors, whereas the neuronal agreement varies by individual and region; within the Zhou et al. cohort, the two other donors also disagree with each other on neurons (rho 0.17). Oligodendrocyte and OPC correlations are low in every arm (rho ≤ 0.17 and ≤ 0.29), and the inhibitory-neuron arms without donor 861 yield no gene significant in both cohorts. These are consistency measures, not replication, and the arms are not compared by test. The statistics and the within-cohort donor agreement are in Supplementary Data 9, sheet Donor arms; the twelve-comparison summary is in Supplementary Data 9, sheet Direction consistency. The control groups of the two studies are independent.

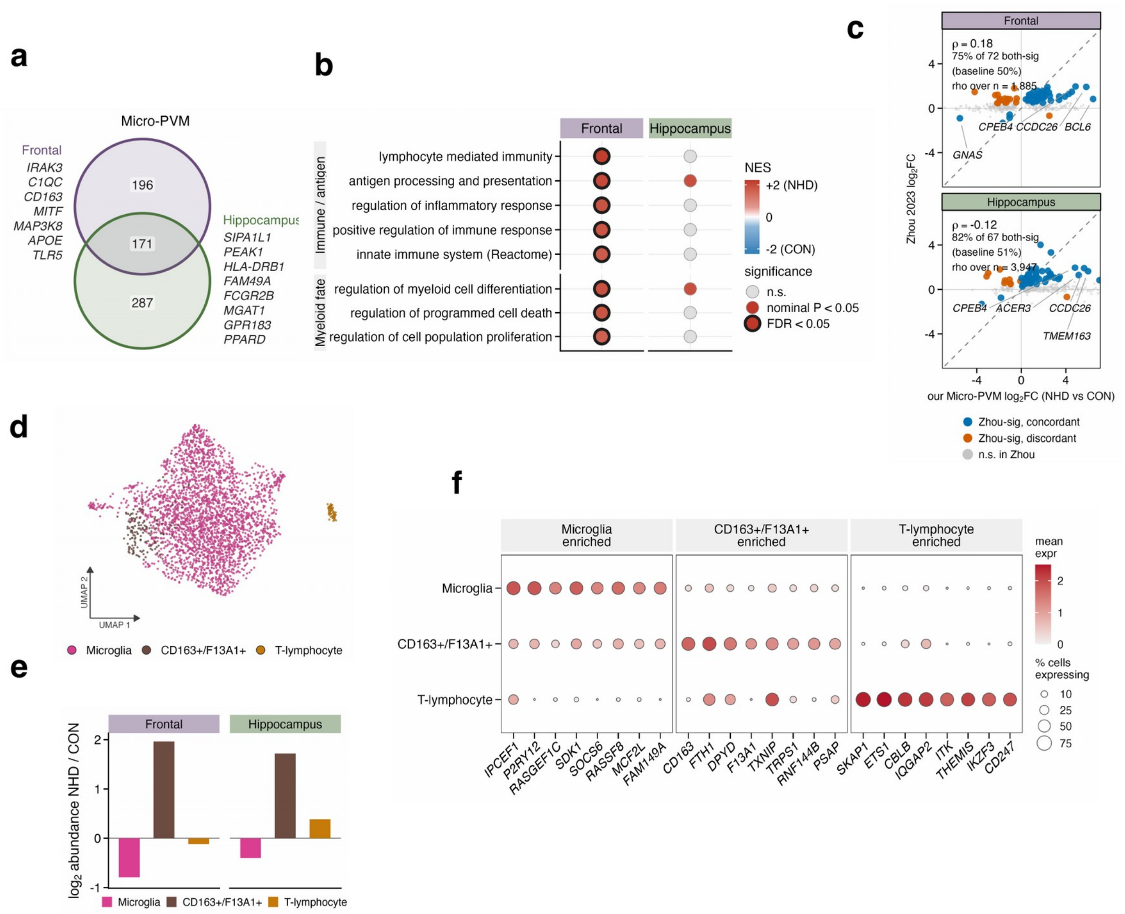


**Supplementary Fig. 3 Myeloid subpopulations, pathway enrichment, and cross-cohort direction consistency of the microglial response. a** Micro-PVM discovery features shared between regions on the unfiltered discovery set (196 prefrontal-only, 171 in both, and 287 hippocampus-only; 163 of the 171 agree in direction), with representative genes. The filtered counts in Fig. 1c are the same analysis after the artifact filter. **b** Pathway enrichment of NHD versus control Micro-PVM (GO biological process and Reactome). fgsea was run on the signed per-nucleus log2 fold change, restricted to genes detected in ≥ 10% of nuclei in both conditions and free of ambient neuronal markers, with Benjamini-Hochberg correction within region, gene sets of ≥ 15 genes, and terms sharing ≥ 50% of their leading edge collapsed to the member with the best FDR. Dots are of uniform size; fill shows the normalized enrichment score where nominal P < 0.05 (grey otherwise), and a dark ring marks FDR < 0.05. Eight terms were enriched in NHD prefrontal Micro-PVM at FDR < 0.05, led by lymphocyte-mediated immunity (2.19, 0.006; leading edge *FCGR3A*, *HLA-DRA*, *C1QA/B/C*, *B2M*, *FCGR1A*, *HLA-DRB1*, *CD74*, and *TYROBP*, which are complement, MHC class I and II, Fc-receptor, and adaptor genes rather than lymphocyte genes), antigen processing and presentation (2.03, 0.047), regulation of the inflammatory response (1.86, 0.047), and the Reactome innate immune system (1.71, 0.028). In hippocampus, which has a single NHD library, two terms reached nominal P < 0.05 and none reached FDR < 0.05, so the hippocampal result is interpreted for direction only. No term was lost in NHD in either region. **c** Agreement in direction with the Zhou et al. cohort (three NHD donors: ours, one further c.2T>C homozygote, and one homozygous *TYROBP* c.141delG carrier; eleven controls). Our Micro-PVM log2 fold change is plotted against theirs: blue, same direction in both cohorts; vermilion, opposite; grey, not significant in Zhou et al. Of the genes significant in both cohorts, 75% of 72 agree in prefrontal cortex and 82% of 67 in hippocampus, against empirical baselines of 50.0% and 50.9% over 1,885 and 3,947 shared genes (binomial P = $1 \times 10^{-5}$ and $1 \times 10^{-7}$). Spearman rho over the shared set is +0.18 and −0.12; the hippocampal value is anti-correlated genome-wide (P = $4 \times 10^{-14}$, stable to leave-one-out), so the hippocampal arm is interpreted as agreement on the strongest shared genes only. The regions are not matched between cohorts. **d** Re-clustering of the immune subset (3,046 nuclei): microglia (2,824), a $CD163^{+}/F13A1^{+}$ population (162), and T lymphocytes (60). The T lymphocytes form a discrete island, excluded from the myeloid analyses, which is drawn shifted toward the main cloud with its geometry unchanged. **e** log2 abundance ratio (NHD/CON, per 10,000 nuclei) by class and region: microglia −0.79 (prefrontal) and −0.40 (hippocampus); $CD163^{+}/F13A1^{+}$ +1.97 and +1.72, on small counts (19 to 47 nuclei in prefrontal cortex and 53 to 43 in hippocampus, a relative increase against the smaller NHD denominator); T lymphocytes −0.12 and +0.39. These are effect sizes without tests. **f** Class-distinguishing markers: the $CD163^{+}/F13A1^{+}$ block is only partly private, whereas the T-lymphocyte block is unambiguous. Dot size, percentage of nuclei expressing; fill, mean log-normalized expression. The origin of the $CD163^{+}/F13A1^{+}$ population is unresolved, and no perivascular or peripheral origin is asserted. Source data: Supplementary Data 3, 5, and 9.

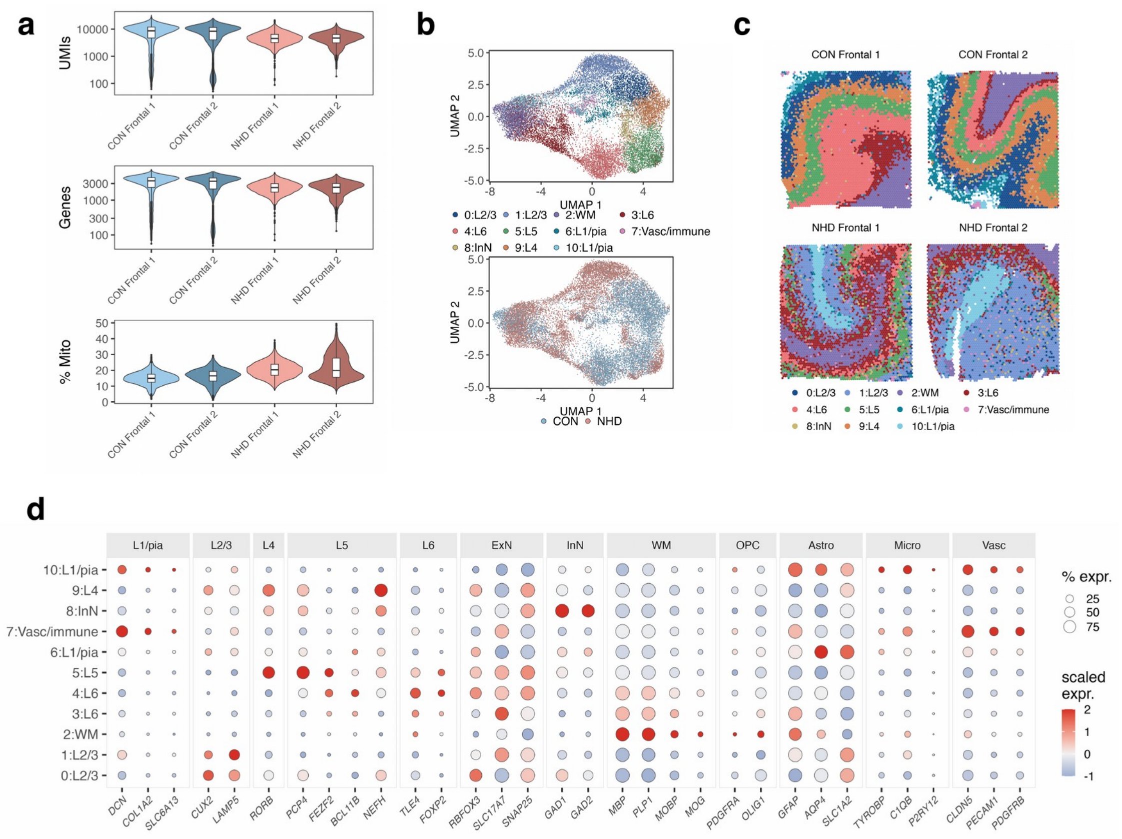


**Supplementary Fig. 4 Visium quality control and spatial-domain assignment.** Spatial transcriptomics profiled prefrontal cortex from one control and one NHD donor, with two sections each. A total of 18,765 spots (CON 9,359, NHD 9,406) cleared a condition-blind floor of 50 UMIs and 25 genes, with no mitochondrial ceiling; the percentage of mitochondrial reads was regressed out before clustering. **a** Per-spot UMIs, detected genes, and mitochondrial fraction by section: median 8,487 UMIs and 3,564 genes in the control sections, against 4,598 and 2,327 in NHD, with a higher mitochondrial fraction in NHD (20.2% against 15.7%). **b** Harmony integration resolved nine clusters. The control upper-cortex cluster was subclustered into a *CUX2*/*LAMP5*-high L2/3 band and a *RORB*-high L4 band, and the BANKSY-guided L1/pia assignment is kept as a separate cluster id, giving eleven cluster ids (upper; 0–10) that collapse to eight named domains recovered in both conditions (lower). Occupancy is skewed by condition (4% to 95% NHD spots across clusters, 4% to 79% across domains) and decreases with depth (Spearman rho −0.57 over the eight domains). **c** Domains on tissue in the frame of Fig. 6, following the pial surface and the grey–white border of each section. **d** Marker basis of the domain labels: astrocytic *GFAP*, *AQP4*, and *SLC1A2* at the tissue edge and leptomeningeal *DCN*, *COL1A2*, and *SLC6A13* (L1/pia); *CUX2* and *LAMP5* (L2/3); *RORB* (L4); *PCP4*, *FEZF2*, *BCL11B*, and *NEFH* (L5); *TLE4* and *FOXP2* (L6); *GAD1* and *GAD2* (InN); *MBP*, *PLP1*, *MOBP*, and *MOG* (WM); and *CLDN5*, *PECAM1*, and *PDGFRB* with *TYROBP*, *C1QB*, and *P2RY12* (vascular/immune). Fill, z-scored mean capped between −1 and +2; dot size, percentage of spots expressing. The sections are technical replicates of one donor per condition, so the panels are descriptive, with no statistical comparison. Source data: Supplementary Data 11 and 2.

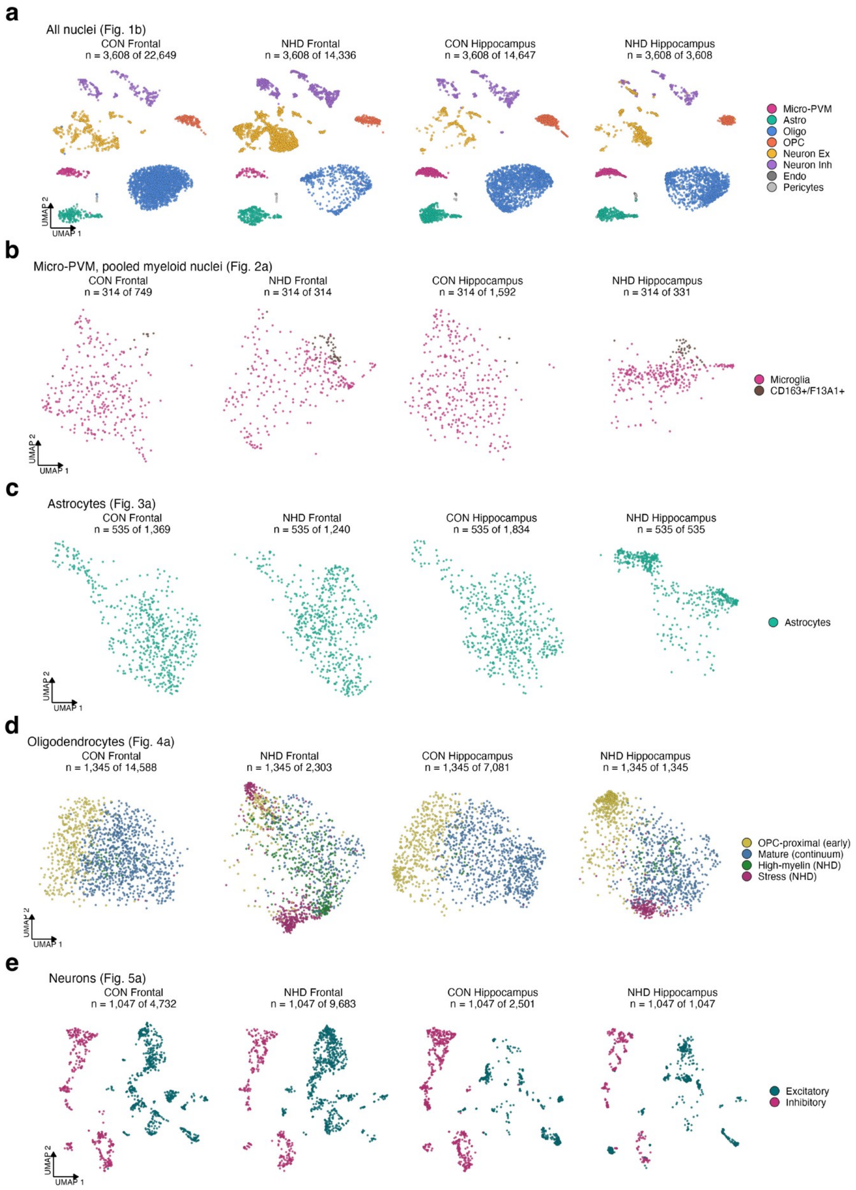


**Supplementary Fig. 5 Each main-figure embedding drawn separately by condition and region.** The five embeddings of the main figures on their own joint coordinates in four panels, CON frontal, NHD frontal, CON hippocampus, and NHD hippocampus, with identical axis limits, point size, and color within each row: **a** all nuclei by cell class (Fig. 1b); **b** myeloid nuclei by class, T lymphocytes excluded (Fig. 2a); **c** astrocytes (Fig. 3a), cropped as in that panel; **d** oligodendrocytes by continuum state (Fig. 4a); **e** neurons by class (Fig. 5a). Each panel is down-sampled without replacement to the smallest of its four groups (seed 42), so a denser cloud is a different distribution rather than more nuclei; each title gives n drawn of n total. All eight classes are present in every group (**a**), the $CD163^{+}/F13A1^{+}$ nuclei in all four (**b**), excitatory and inhibitory territories are shared (**e**), and the two NHD oligodendrocyte states are occupied almost entirely by NHD nuclei, CON contributing 2.9% and 0.2% of the frontal lineage (**d**). One donor per condition; the NHD hippocampus is a single sample. Nuclei outside a cropped frame are retained in the embedding but not drawn.

### Supplementary tables

**Supplementary Table S1** Gene-level criteria for the TREM2-independent (DAM-1) and TREM2-dependent (DAM-2) arms in Micro-PVM, from Supplementary Data 3. A discovery feature requires Bonferroni-adjusted $P < 0.05$, $|\text{Cliff's delta}| \geq 0.15$ and detection $\geq 10\%$ in one condition. “Not returned” means the gene did not pass the test’s prefiltering criteria in that region.

| Arm | Gene | Region | Cliff’s delta | Detection NHD / CON (%) | Adjusted P | Effect-size criterion | Detection criterion | Significance criterion |
|---|---|---|---|---|---|---|---|---|
| DAM-1 | *P2RY12* | Prefrontal | −0.39 | 33 / 66 | $3.4 \times 10^{-25}$ | met | met | met |
| DAM-1 | *P2RY12* | Hippocampus | −0.46 | 41 / 66 | $6.1 \times 10^{-27}$ | met | met | met |
| DAM-1 | *APOE* | Prefrontal | +0.18 | 39 / 23 | $6.7 \times 10^{-5}$ | met | met | met |
| DAM-1 | *APOE* | Hippocampus | +0.23 | 49 / 22 | 1 | met | met | not met |
| DAM-2 | *CLEC7A* | Prefrontal | +0.09 | 19 / 10 | 1 | not met | met | not met |
| DAM-2 | *CLEC7A* | Hippocampus | +0.21 | 36 / 12 | 1 | met | met | not met |
| DAM-2 | *ITGAX* | Prefrontal | −0.02 | 26 / 28 | 1 | not met | met | not met |
| DAM-2 | *ITGAX* | Hippocampus | +0.15 | 51 / 28 | 0.16 | met | met | not met |
| DAM-2 | *GPNMB* | Prefrontal | — | not returned | — | — | — | — |
| DAM-2 | *GPNMB* | Hippocampus | +0.13 | 14.5 / 1.0 | 1 | not met | met | not met |
| DAM-2 | *TREM2* | Prefrontal | +0.02 | 11 / 8 | 1 | not met | met | not met |
| DAM-2 | *TREM2* | Hippocampus | +0.148 | 22 / 6 | 1 | not met | met | not met |
| DAM-2 | *PPARG* | Prefrontal | — | not returned | — | — | — | — |
| DAM-2 | *PPARG* | Hippocampus | −0.06 | 7 / 12 | 0.51 | not met | met | not met |
| DAM-2 | *LPL*, *CST7*, *LGALS3* | both | — | not returned | — | — | — | — |

### Supplementary Data legends

**Supplementary Data 1** Data dictionary: the columns of every sheet of every Supplementary Data workbook, with the workbook, sheet, figures backed and source script of each table, and the analysis-wide conventions. Each workbook opens on a README sheet that defines its columns.

**Supplementary Data 2** Samples, nuclei and annotation. Sheets: QC per sample (sequencing and quality-control metrics for the seven single-nucleus samples); Nucleus counts (post-QC nuclei per cell type, region and condition); Frontal subclasses (cortical neuron subclass counts per condition, 15 subclasses); Hippocampal subtypes (subtype assignment per hippocampal neuronal nucleus); Myeloid selection (the myeloid nucleus selection chain per condition and region).

**Supplementary Data 3** One data sheet: per-nucleus MAST differential expression, NHD versus control, per cell type and region and per frontal neuronal subclass, with Bonferroni-adjusted P, Cliff's delta, detection and the discovery gate.

**Supplementary Data 4** One data sheet: pseudobulk DESeq2 differential expression per cell type and region (the confirmation tier; hippocampal P values withheld).

**Supplementary Data 5** Gene-set enrichment. Sheets: Data (fgsea on genes ranked by trimmed log2 fold change, per cell type and region, with nucleus-count power tiers); Micro PVM detection gated (the detection-gated microglial enrichment behind Supplementary Fig. 3b).

**Supplementary Data 6** Gene sets and markers. Sheets: Signatures (every curated signature and program gene set used for module scoring and dot plots, with symbol-mapping provenance and any exclusion); Cell-type markers (top ten protein-coding positive markers per annotated cell type).

**Supplementary Data 7** Program effect sizes and sensitivity. Sheets: Program deltas and By program (Cliff's delta, NHD relative to control, with Benjamini-Hochberg q and the effect gate for every program of Figs 2d, 3e, 4d and 5d, long and wide); Neuron scores per nucleus (the eight neuronal program scores per nucleus); Unresolved sensitivity (hippocampal excitatory-neuron program deltas with and without the unresolved cluster); RIN per library and RIN gap (per-sample mean excitatory-neuron program scores beside RNA integrity, and the per-region integrity gap against program loss).

**Supplementary Data 8** Microglia. Sheets: TF activity Fig2h (CollecTRI regulon activity across the myeloid nuclei behind Fig. 2h, with the six-criterion admission gate and the 750-UMI down-sampling sensitivity arm); Pooled vs microglia-only (effect sizes for eight activation genes on the pooled Micro-PVM class and on microglia alone).

**Supplementary Data 9** Cross-cohort contrast with Zhou et al. Sheets: Direction consistency (agreement in direction per cell type and region against the empirical rate of chance agreement); Donor arms (the same statistic recomputed with Zhou's NHD side restricted by donor, with and without the shared donor); Within Zhou donors (donor-to-donor agreement within the Zhou cohort); TYROBP per donor (TYROBP, TREM2 and CSF1R detection in the microglia of each Zhou et al. donor, with median sequencing depth).

**Supplementary Data 10** Sequencing-depth diagnostics. Sheets: A relative RNA content (per-compartment NHD-to-control depth ratios); B threshold sweep (the UMI-floor sweep); C artifact by treatment (the depth-only artifact by normalization); D condition balance; E neuron module artifact (the neuronal-program depth bound); F myeloid condition depth (the hippocampal microglial RNA-content comparison and the down-sampling sensitivity).

**Supplementary Data 11** Visium spatial transcriptomics, in the order of Fig. 6. Sheets: Spot annotation (every analysed spot with section, condition, cluster, domain, compartment, tissue-call tier and the layer I/pia flags) and Layer IV order (descriptive per-section statistics of the L4 IT marker signature); Programs per domain, Genes by band and Gene sets Visium (program expression per domain and condition, the per-gene band means and log2 ratios of the Fig. 6d heatmap, and the gene membership of every program scored on tissue); Microglia proximity (Cliff's delta between microglia-rich and microglia-poor spots per program, compartment and section, with block-bootstrap intervals and the toroidal-shift null); Composition per domain (cell2location mean proportions per domain and condition with log2 ratios and block-bootstrap intervals).

**Supplementary Data 12** Source data for the cell-cell communication and composition panels. Sheets: Compensatory routes (per-gene control and NHD detection of the receptor, adaptor and kinase genes of Fig. 2c, with the depth-adjusted change in percentage points and the detection floor); Chord pathway totals (summed CellChat communication probability per sender, receiver and pathway behind the chord diagrams of Figs. 3g, 3h and 5e, with each ribbon's share); LR microglia to glia (ligand-receptor communication probabilities per condition and region behind Figs. 3g and 4j, with permutation P values at the 0.01 resolution of 100 permutations); LR astrocyte to neuron (the pairs of Fig. 3h, with the raw-count detection, differential-expression delta and discovery status of each receptor in the receiving class); LR microglia to neuron (the pairs of Fig. 5e per region and neuronal class); LR microglia to neuron subtype (mean expression and detection of each receptor per neuronal subtype, region and condition behind Fig. 5g); Oligo state composition (nuclei and fraction per oligodendrocyte state, region and condition behind Fig. 4h).
